# Introgressed *Manihot glaziovii* Alleles in Modern Cassava Germplasm Benefit Important Traits and Are Under Balancing Selection

**DOI:** 10.1101/624114

**Authors:** Marnin D. Wolfe, Guillaume J. Bauchet, Ariel W. Chan, Roberto Lozano, Punna Ramu, Chiedozie Egesi, Robert Kawuki, Peter Kulakow, Ismail Rabbi, Jean-Luc Jannink

## Abstract

1.

Introgression of alleles from wild relatives has often been adaptive, usually for disease resistance, in plant breeding. However, the significance of historical hybridization events in modern breeding is often not clear. Cassava (*Manihot esculenta*) is among the most important staple foods in the world, sustaining hundreds of millions of people in the tropics, especially in sub-Saharan Africa. Widespread genotyping makes cassava a model for clonally-propagated root and tuber crops in the developing world and provides an opportunity to study the modern benefits and consequences of historical introgression. We detected large introgressed *M. glaziovii* genome-segments in a collection of 2742 modern cassava landraces and elite germplasm, the legacy of 1930’s era breeding to combat epidemics disease. African landraces and improved varieties were on average 3.8% (max 13.6%) introgressed. Introgressions accounted for significant (mean 20%, max 56%) portion of the heritability of tested traits *M. glaziovii* alleles on the distal 10Mb of chr. 1 increased dry matter and root number. On chr. 4, introgressed alleles in a 20Mb region improved harvest index and brown streak disease tolerance. Three cycles of selection initially doubled the introgression frequency on chr. 1. Later stage variety trials selectively excluded homozygotes which indicates a heterozygous advantage. We show that maintaining large recombination-suppressed introgressions in the heterozygous state allows the accumulation of deleterious mutations. We conclude that targeted recombination of introgression segments would therefore increase the efficiency of cassava breeding by allowing simultaneous fixation of beneficial alleles and purging of genetic load.

**Significance Statement:** Crosses to wild relatives have often been adaptive for crop breeding, but their modern importance is usually poorly understood. Cassava (*Manihot esculenta*) is an important staple crop, feeding hundreds of millions in the developing world, and is a model for vegetatively-propagated non-inbred crops. In the 1930’s, crossing to *M. glaziovii* averted mosaic disease epidemic in Africa. We reveal that large genome segments, the legacy of those crosses, benefit a number of traits including yield in modern cassava and are consistently favored during selection. Elite cultivars are almost exclusively heterozygous for wild alleles; homozygotes are rejected during early stage trials, suggesting inbreeding depression. More recombination around beneficial wild alleles will allow purging of genetic load and increase genetic gain in cassava.

## 2. Introduction

Interspecific hybridization has provided an important source of adaptive genetic variation during the evolution in many organisms including humans (1, 2), cattle (3) and maize (4). Indeed introgression between many crops and their undomesticated relatives has occurred in both directions (5), naturally in farmers fields and deliberately by plant breeders (6–9). Introgression can also have serious population genetic consequences including genomic inversions and other structural variations, suppression of recombination and segregation distortion, inbreeding depression and hybrid sterility (10–12). Despite many individual examples, the consequences of historical introgressions both positive and negative, especially at the quantitative genetic level, is rarely simultaneously understood.

Cassava (*Manihot esculenta*) is among the most important staple foods in the world, sustaining hundreds of millions of people in the tropics, especially in sub-Saharan Africa (http://faostat.fao.org). Cassava is a clonally-propagated staple food crop, grown throughout the tropics for its starchy storage roots. In recent years, cassava has emerged from orphan-crop status to a model for plant breeding in the developing world generally, especially for outbreeding non-cereals and vegetatively-propagated root and tuber crops (13–16).

The history of cassava breeding includes periodic tapping of wild con-generic relatives as sources of useful genetic variation (7, 17). In the early 20th century, cassava production in Africa faced a grave threat in the form of mosaic disease, a geminivirus caused, insect-vectored pathogen. Records indicate that an initial worldwide search for resistant cultivated cassava was conducted ((18–21)). Failing to find native resistance, breeders at the Amani research station in Tanzania introgressed resistance from the Ceara rubber tree (*Manihot glaziovii* Muell. Arg.) (18–22).

Three backcrosses of hybrids to *M*. *esculenta* produced acceptable levels of resistance and storage root yield (18, 22) leading to the distribution of mosaic tolerant varieties to farmers in the local area of Amani (18, 23) and the eventual end of the first mosaic disease epidemics by the 1940’s (18, 23). Descendants of these original hybrids became key founders of modern breeding germplasm (21, 22, 24). The Amani derived lines have been identified as important sources of resistance against cassava mosaic disease (CMD) (25, 26), brown streak disease (CBSD) (27) and bacterial blight (28).

Large genome-segments derived from *M. glaziovii* were recently discovered in a sample of African genotypes, suggesting that historical introgressions remain important today (29). Several other studies have identified QTL in these regions, leading us to hypothesize that *M. glaziovii* alleles confer CBSD resistance (29–31) and possibly increased storage root dry matter content (32).

Widespread genotyping for genomic selection (GS) in African (www.nextgencassava.org) cassava breeding makes cassava a model for root and tuber crops in the developing world and provides an opportunity to study the modern benefits and consequences of historical introgression. We leveraged publicly available data (www.cassavabase.org) from more than 2742 breeding lines, land races and local varieties, with both field phenotypes and genome-wide marker records (16) as well as whole-genome sequences (33). First, we investigated the legacy of *M. glaziovii* introgression by determining its extent in the germplasm and the associated population structure. We then employed a combination of genetic variance partitioning, genome-wide association analysis and genomic prediction to quantify the location, effects and overall importance of introgressed alleles for key cassava traits and thus for cassava breeding. Finally, we study three generations of genomic selection progenies to understand the role of introgressions in modern cassava breeding.

## Results

### Introgression-associated population structure

In order to detect introgressed *M. glaziovii* genome segments in cultivated cassava samples, we defined introgression diagnostic markers (IDMs) across the genome using an approach similar to that of Bredeson et al. (29). We contrasted allele frequencies between a panel of *M. glaziovii* and a set of “non-introgressed” *M. esculenta*. The cassava HapMapII, a 30X WGS diversity panel (33), includes eight accessions identifiable as *M. glaziovii*. Defining a *M. esculenta* panel required additional analysis, as some of our samples are themselves introgressed. We defined 1000 SNP-window specific sets of 10 *M. esculenta*, which were least likely to be introgressed in that window because they were the most genetically distant from the *M. glaziovii* samples (Figs. S1-2).

A jump in the variability of genetic distance from *M. glaziovii* in HapMapII clones occurred on chromosome 1 from Mb 25 to the end of the chromosome (Fig. S2). This area corresponds to a region shown previously to be segregating for *M. glaziovii* introgressions (29). Ultimately, we considered only the 120,990 sites intersecting between the HapMapII and the broader genotyping-by-sequencing (GBS) dataset, we intended to analyze. From this set, we identified 38000 IDMs that were either fixed for opposite alleles (N=20,681) or fixed in the *M. esculenta* reference panel, but polymorphic amongst the *M. glaziovii* (N=17,319). At each IDM locus, therefore, we could identify an allele that was putatively derived from *M. glaziovii* and would be diagnostic of introgression if found in a cultivated cassava genome (Table S1). The IDMs were distributed similarly across the genome compared to the rest of the markers in our dataset (Fig. S3).

Principal components analysis (PCA) on the IDM markers revealed a pattern of relatedness in introgression regions (Fig. 1A) that is distinct from that of the rest of the genome (Fig. 1B) or overall (Fig. 1C). We coded the dosages for the IDMs to count the *M. glaziovii* diagnostic allele. The resulting loadings (eigenvector coefficients) for markers on PC1 (21% variance explained) are strongest for IDMs on the last 10 Mb of chr. 1, while the strongest loadings on PC2 (9%) are at IDMs spanning the majority of chr. 4 (Fig. 1D).

**Fig. 1.**
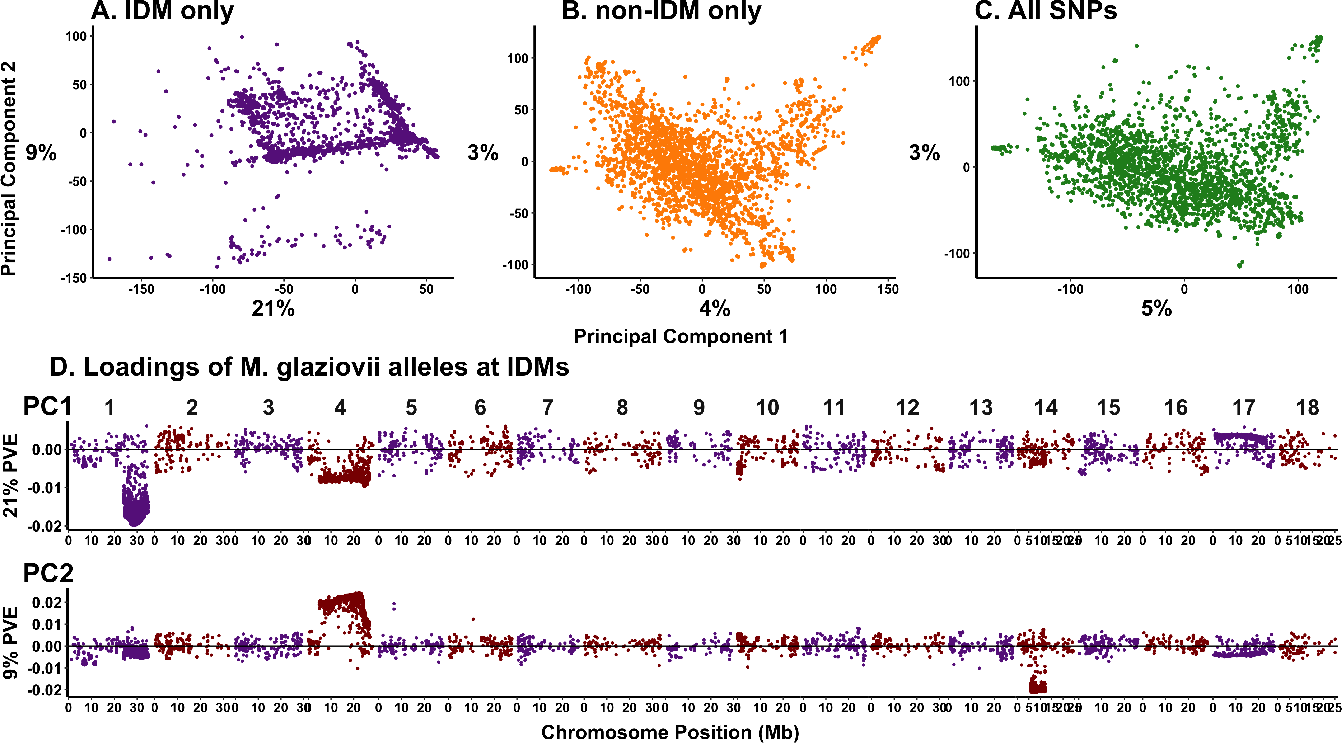
Introgression regions capture distinct population structure. Shown are scatterplots of the PC scores for PC1 and PC2 from 3 PCAs using 3 sets of markers: all together (C, includes “tag”-IDM), non-IDM only (B), IDM only (A, excludes “tag”-IDM). D. The loadings or eigenvector coefficients from the IDM-only PCA are shown plotted against their genomic coordinates for the first two principal components (vertical panels).

### Introgression frequency divergence among populations

The genome-wide proportion of *M. glaziovii* alleles per clone ranged from 1.3% to 13.6% (mean 3.8%) among the African breeding germplasm as a whole (GG+LG+NR+UG; Fig. 2B; Tables S2-3). On a genome-wide basis, there are not large differences among populations in the mean frequency of introgressed alleles. The breeding populations GG (4.2%), NR and UG (4.1%) have similar levels of introgression, while the L. American collection was the least introgressed (1.8%) and the local germplasm (LG, 3%) were intermediate. We note that in the CIAT collection, the Brazilian accession BRA534 appears to be an outlier with 34% *M. glaziovii* alleles. We excluded BRA534 when comparing CIAT to other populations.

**Fig. 2.**
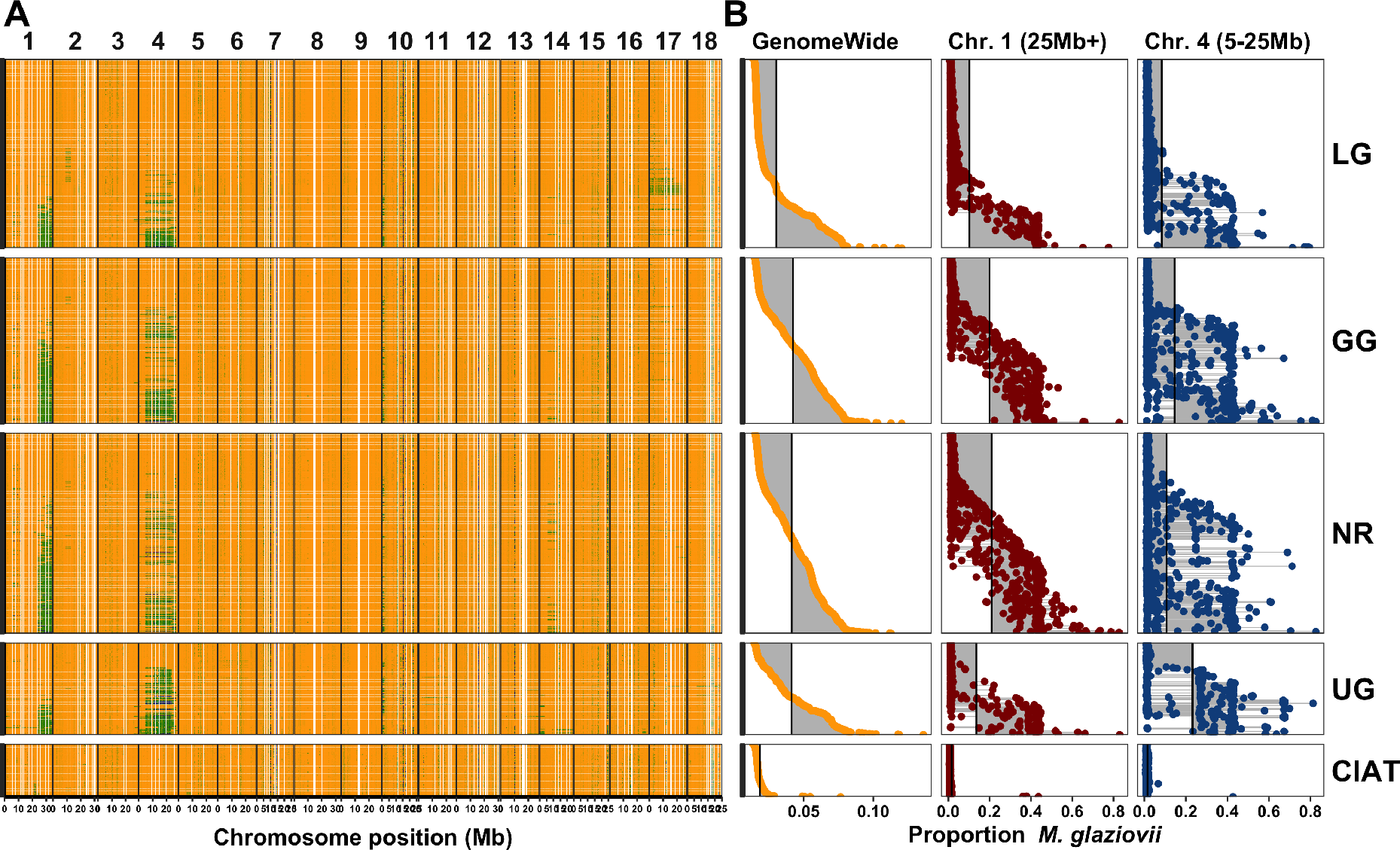
Comparison of introgression among populations. The mean *M. glaziovii* allele dosage at IDMs in 250Kb windows across the genome is depicted on the left (A). Physical position on each chromosome is depicted in megabases (Mb) along the x-axis. Colors range from orange (0 M.g. alleles), to green (1 M.g. allele), to dark blue (2 M.g. alleles). The per individual proportion of *M. glaziovii* alleles at IDM is summarized on the right (B). Proportions were calculated as the sum (per clone) of the dosages at IDMs divided by two times the number of IDMs. The proportions in B were computed either with all IDMs (GenomeWide, left column), at IDMs on chr. 1 >25 Mb (middle column), at IDMs on chr. 4 from 5 to 25 Mb (right column). The populations can be compared in B by looking down the columns and using the vertical lines, which represent the mean values for that group and region, as a visual aid. For both A and B, each row (y-axis) is an individual cassava clone and the vertical panels represent five populations: IITA local germplasm (LG), IITA Genetic Gain (GG), NRCRI (NR), NaCRRI (UG) and the L. American collection (CIAT). Rows (clones) are aligned across A and B and sorted within population based on the genome-wide proportion *M. glaziovii* (left column of B).

The largest introgressions detected were apparently contiguous segments of chr. 1 approx. 25Mb to the end (10Mb total) and chr. 4 from 5Mb to 25Mb (Fig. 2A). When we isolate the introgressions on chrs. 1 and 4, which appear to be the same as were previously identified (29), we observe more striking differences. The frequency of the chr. 1 segment was on average greater in the W. African breeding germplasm GG (0.2) and NR (0.21) than in the E. African population UG (0.14). In contrast, the introgression on chr. 4 was more common in UG (0.23) compared to GG (0.15) or NR (0.11). Samples from the IITA local germplasm collection (LG) were less likely to contain introgressions on either chrs. 1 (0.10) or 4 (0.08) and the L. American samples from CIAT showed almost no evidence of introgression (<0.02) on both chrs. 1 and 4 (Fig. 2B, Tables S2-3).

### Ongoing selection for *M. glaziovii* alleles

We compared the introgressions detected in local germplasm and landraces of cassava (LG) to IITA improved varieties (GG) and three successive generations of genomic selection progeny (C1, C2, and C3), which descend from parents selected initially from the GG. The most notable changes we observed were on chrs. 1 and 4 (Fig. 3A). Genome-wide, the average proportion of *M. glaziovii* alleles per individual increased from 0.03 in LG to 0.042 in GG and then more than doubled in the GS progeny with C1 at 0.095, C2 and C3 at 0.12 (Fig. 3B; Table S2-3). Most of this change was due to increases on chr. 1, which rose from 0.1 in the LG to 0.2 in the GG and maxed out at 0.34 in the C3. In contrast, the chr. 4 region appears to have stayed steady around 15% from GG through C2 and even slightly decreases from C2 to C3 (Fig. 3B, Tables S2-3).

**Fig. 3.**
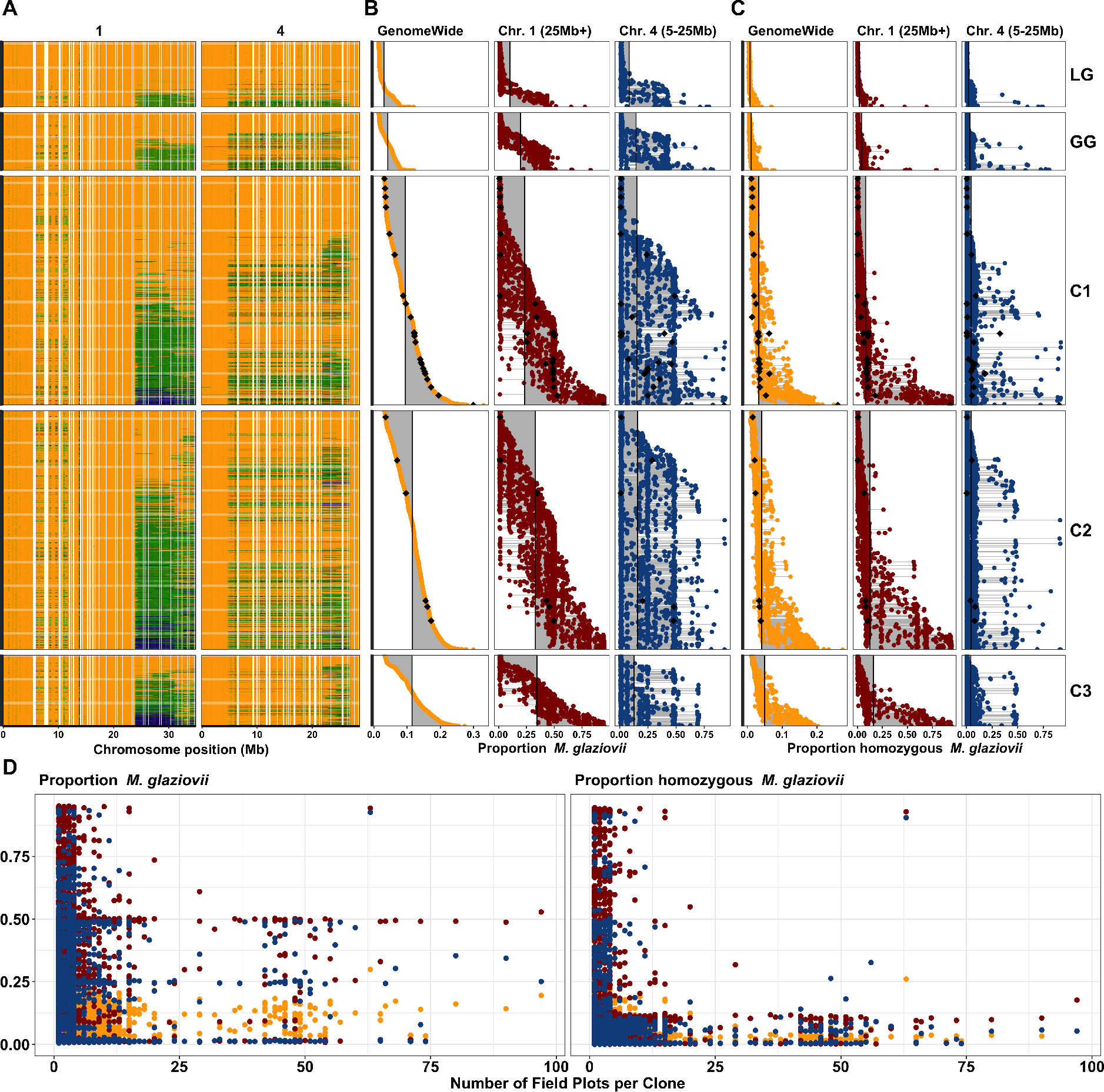
The effects of (genomic) selection on *M. glaziovii* introgressions. The mean *M. glaziovii* allele dosage at IDMs in 250Kb windows on chromosomes 1 and 4 is depicted on the top left (A). Physical position on each chromosome is depicted in megabases (Mb) along the x-axis. Colors range from orange (0 M.g. alleles), to green (1 M.g. allele), to dark blue (2 M.g. alleles). The top middle panel (B) shows the per individual proportion and the top right panel (C) shows the rate of homozygosity for *M. glaziovii* alleles at IDM. Proportions for B were calculated as the sum (per clone) of the dosages at IDMs divided by two times the number of IDMs. The proportions for C were simply the proportion (per clone) out of the total number of IDMs with a dosage equal to two. The proportions in B and C were computed either with all IDMs (GenomeWide, left column), at IDMs on chr. 1 >25 Mb (middle column), at IDMs on chr. 4 from 5 to 25 Mb (right column). The populations can be compared in B and C by looking down the columns and using the vertical lines, which represent the mean values for that group and region, as a visual aid. For A through C, each row (y-axis) is an individual cassava clone and the vertical panels represent five populations: IITA Genetic Gain (GG) and three successive generations of genomic selection progeny (C1, C2 and C3), descended originally from GG. Rows (clones) are aligned across A-C and sorted within population based on the genome-wide proportion *M. glaziovii* (left column of B). At the bottom, D shows how the introgression frequency and homozygosity rate per individual (y-axis) for the C1, C2 and C3 relates to the cumulative number of field plots planted (as of Jan. 2019) per clone (x-axis). The number of field plots per clone is meant as a proxy representing the level of advancement through variety development stages of the breeding process. For illustrative purposes, we highlight C1, C2 and C3 clones with >50 field plots in panels B and C with black diamonds. For D as in B and C, we break down the proportions in D by region and use the same color coding: genome-wide (orange), chr. 1 region (dark red), chr. 4 region (dark blue).

Most introgressed LG and GG were heterozygous for *M. glaziovii* haplotypes, with a mean homozygosity rate of only 1% genome-wide (Fig. 2A, Fig. 3C). Genomic selection appears to have steadily increased the homozygosity rate on chr. 1 from 4% in the GG to 16% in C3 (Fig. 3A, Fig. 3C, Tables S2-3). The near absence of homozygotes in the elite germplasm (GG) and the gradual increase due to select that we observed, led us to investigate further.

We hypothesized that post-genotyping performance-based selection and advancement through the variety testing process might exclude homozygous clones. We used the cumulative number of field plots planted (according to http://www.cassavabase.org, January 2019) as a metric of the level of advancement each progeny had attained. We found that while heterozygosity for introgressions was acceptable (Fig. 3D, left), homozygous clones were almost completely excluded from later stages (Fig. 3D, right). Of the 30 clones with greater than 50 field plots only one of them appeared to be notably homozygous. For that one clone, both chrs. 1 and 4 were nearly completely homozygous (Fig. 3D, Table S2).

One potential consequence of increasing the frequency of such a large haplotype and maintaining it in a heterozygous state might be the accumulation of deleterious mutations (33). Using a dataset consisting of the LG, GG and C1 with 5.367 million HapMapII SNPs imputed we were able to genotype 9779 putative deleterious mutations of the 22495 identified by (33). From LG to GG, we observed increases in the average per individual genetic load that were larger (34% on chr. 1, 20% on chr. 4) in introgression regions compared to genome-wide (8.7%). Similarly, from GG to C1, genetic load increased, less than between LG and GG, but more in introgression regions (9% for chr. 1 and 4.9% on chr. 4) than genome-wide (2.5%). There was nearly no mean difference between LG and GG in terms of homozygous genetic load. However, there was an increase from GG to C1 and it was also larger in the introgression regions (59% on chr. 1, 15% on chr. 4) than genome-wide (10%) (Tables S2-3).

### Local admixture as confirmation of detected introgressions in HapMapII

We also used HAPMIX (34), a haplotype-based method for local ancestry inference, to detect *M. glaziovii* introgressions in phased WGS HapMapII samples. We found that the HAPMIX and IDM-based methods largely agree (Fig. S4). Although, we note that *M. glaziovii* segments on Chr. 1 tend to appear smaller in the HAPMIX results.

### Heritability accounted for by introgressions

We quantified the proportion of the total genetic variance that is explainable by introgressions segregating in modern cassava germplasm, for nine traits. We compiled data from 68 field trials (42 IITA, 5 NaCRRI, 21 NRCRI) conducted on 2742 genotyped clones in our study populations (Table S5).

To these data, we fit linear mixed-models with two random-genetic effects, kinship measured using IDM markers and kinship by non-IDM markers. The estimated genetic variances partitioned the heritability into two components: one due to introgression regions 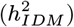and another for the rest-of-the-genome 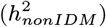. We fit three models: the partitioned model described above (PARTITIONED), a partitioned model with the IDM variance component removed (IDMnull) and a single component, non-partitioned model with kinship from IDM *and* non-IDM markers (ALL). Before fitting these models, we performed two major preliminary analyses.

#### LD between introgressed regions and the rest of the genome

We first investigated the amount of linkage disequilibrium between introgressed and non-introgressed regions. If unaddressed, LD between SNPs in these regions could lead to non-independent estimates of genetic variance when fitting the model described above. The variance arising from introgression regions might then be captured by the non-introgression regions and vice versa (35–37). Using the procedure described in the methods, we reclassified 1413 SNPs, primarily located on chromosomes 1 and 4 (Fig. S7), that were more similar in the kinship they measured to the IDM than to the non-IDM (Figs. S5-7; Tables S1, S4). Redesignating these SNPs as tag-IDM reduced the correlation of IDM and non-IDM kinships from 0.37 to 0.30. We therefore included tag-IDM in the IDM kinship matrices used in all subsequent analyses.

#### Per-trial analyses

The second preliminary analysis we did was to analyze each trial separately, in part in order to check the quality of the data before combining into a larger, multi-trial analysis. Based on a likelihood ratio test, we chose to remove 31 trait-trials that did not show evidence of significant genetic variance (*pLRT* Per-trial analyses. The second preliminary analysis we did was to analyze each trial separately, in part in order to check the quality of the data before combining into a larger, multi-trial analysis. Based on a likelihood ratio test, we chose to remove 31 trait-trials that did not show evidence of significant genetic variance (*pLRT*_*null*_ > 0.05). We removed an additional six trait-trials without any significant genetic variance from marker-estimated covariances. Lastly, 53 more trait-trials were excluded because, based upon the Akaike Information Criterion (AIC), the genomic model fit the data worse than the IID model (Tables S6 & S10)

#### Multi-trial analyses

We combined the remaining trials for each trait (within Institute) to achieve large overall sample sizes (max per Institute: 25924 IITA, 2881 NaCRRI, 6641 NRCRI) and increase the average number of replications per clone (max per Institute: 16.76 IITA, 6.89 NaCRRI, 8.11 NRCRI; Table S11). We fit the three models for each trait and analyzed each breeding program’s data separately.

Ten out of 19 Trait-Institute analyses had significant genetic variance from introgressions. In fact, introgression regions appear important for every trait *except* cassava bacterial blight and mosaic disease severity. In all of these cases, the PARTITIONED model had an AIC more than 2 units smaller than the non-partitioned one. The proportion of the heritability accounted for by significant introgressions was as high as 56% (mean 20%, median 15%, min 3%; Fig. 4, Tables S12-13).

**Fig. 4.**
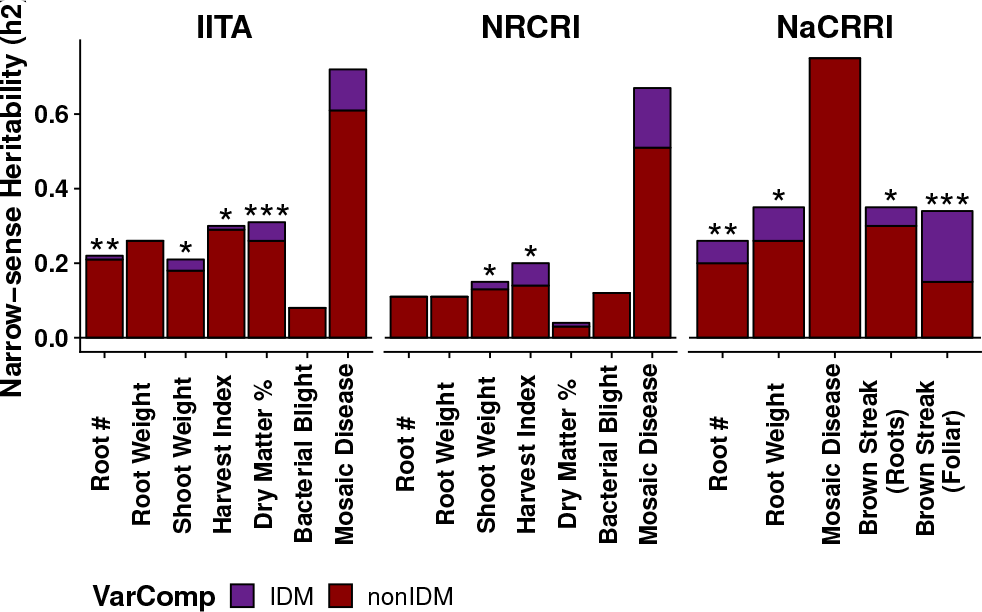
Heritability accounted for by introgressions. The heritability (y-axis) of introgression regions for each trait (x-axis) measured in each breeding program (horizontal panels). Heritability was estimated from partitioned genomic mixed-models and the portion of heritability attributable to introgression regions (purple) vs. the rest of the genome (dark red) is shown. Stars atop bars represent the level of significance in a likelihood ratio test for the significance of the introgression-component (*** p<0.0001, ** p<0.001, * p<0.05).

#### Comparison to random samples

One third of the SNPs in our study were classified as IDMs (including tag-IDMs). We compared the variance explained by our IDM-defined partition, to three random genome partitions of the same size (Table S7). For the random samples, the correlation of GRM’s was >0.99, but was only 0.30 for the IDM-defined partition (Table S8). The IDM-defined partition explained an average of 20% of the total genetic variance, in comparison to 37% for the random partitions, which is closer to proportional with the total number of markers (Fig. S8, Table S12-14).

Most of the cases with significant 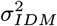 *did not* have significant variance associated with the random samples of equivalent size. In contrast, all three random samples had significant variance for MCMDS in the IITA dataset while the equivalent IDM-defined variance was insignificant.․ For the most part, AIC values indicate the IDM-defined partitions fit at least as well, if not better than the random ones. For only two cases did a random sample appear to fit better than IDM-defined (MCMDS IITA Sample 1, RTNO IITA, Sample 2). In the NaCRRI dataset, IDM-defined partitions fit better than all three random samples for MCMDS and MCBSDS. For the better fitting (compared to random) NaCRRI MCMDS analysis, the variance from the IDM regions was actually zero.

#### Importance of LD

We know from previous studies in cassava, and confirm here (Fig. S9, Table S21), that the introgression regions on chromosomes 1 and 4 are characterized by strong, relatively long-range LD (29, 32). We computed the cumulative genetic size in centimorgans in one megabase windows along each chromosome (Table S21). We found that recombination was 14% and 71% less than the rest of the genome in the introgressions on chrs. 1 (25Mb+) and 4 (5-25Mb), respectively. We used the LDAK method (35) to weight SNPs contributions to kinship matrices (GRMs) in order to correct for variability in tagging of causal mutations due to LD (35, 37) (Figs S10-11, Table S9). The key result of LD adjustment was a mean decrease of 7.4% of the proportion 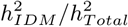 (max decrease −41.6% for CBSDRS, max increase 6.6% for RTNO), among cases where at least one of the models had significant *LRT*_*IDM*_ (Figs. S12-13, Table S17). We noted that the off-diagonals of the LDAK adjusted IDM and non-IDM GRMs were more correlated to each other (0.65) than the unadjusted pair (0.3; Table S16). The IDM matrix was altered most by LDAK adjustment, with off-diagonal correlation to the unadjusted IDM matrix of 0.45 compared to 0.89 for the non-IDMs (Table S15). In all, 12 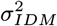 were significant either before, after or both before and after LD adjustment. Of these 12, there were two where LD adjustment made the *LRT*_*IDM*_ significant and three where it became insignificant.

### *M. glaziovii*-associated QTL

We identified quantitative trait loci (QTL) attributable to *M. glaziovii* alleles using mixed linear model GWAS on two types of predictors. The first GWAS was on the SNP markers themselves and the second was on the mean dosage of *M. glaziovii* alleles in 250Kb windows (*DoseGlaz*), described in more detail in METHODS. There were bonferroni-significant IDM and/or *DoseGlaz* for all traits except bacterial blight (MCBBS) (Fig. 5; Figs. S14, S16).

**Fig. 5.**
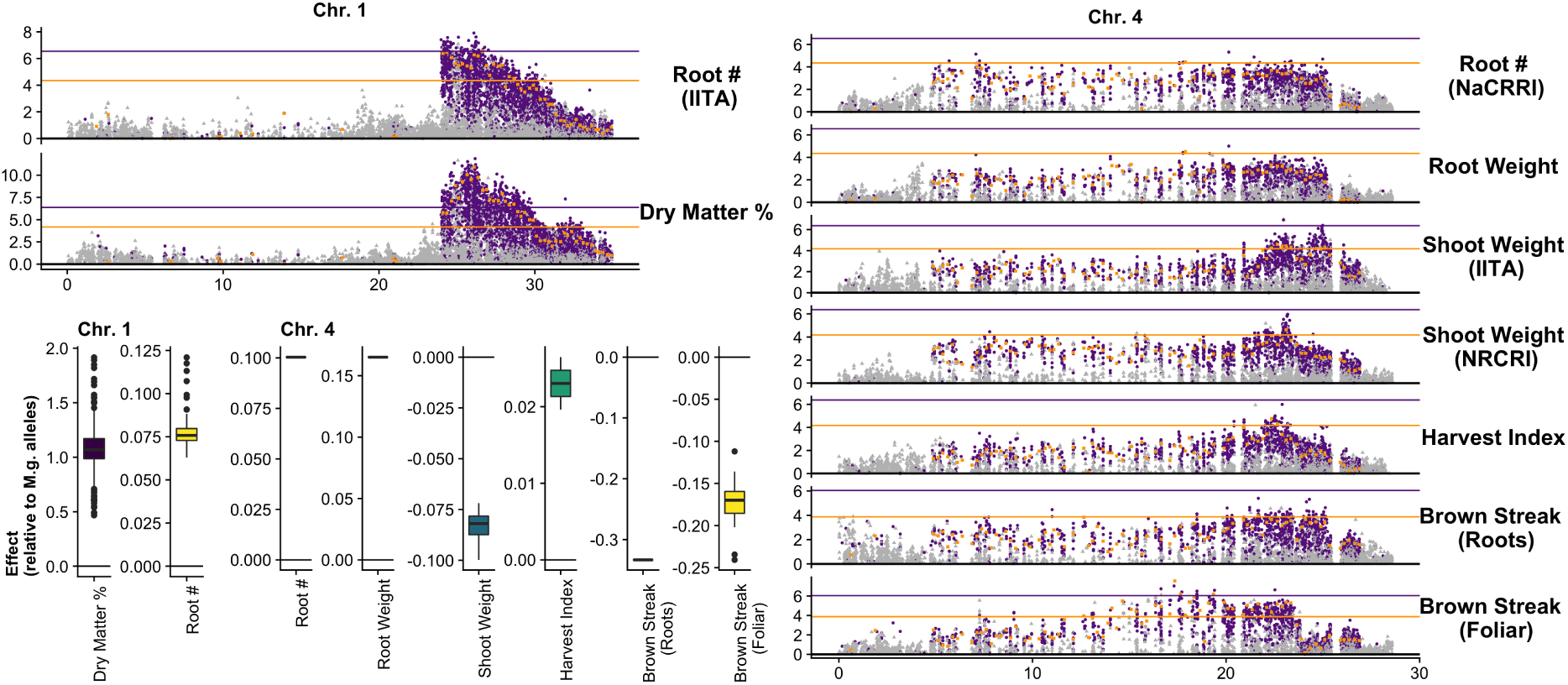
Significant introgression-trait associations. Manhattan plots summarizing genome-wide associations for traits with significant introgression-trait associates on chromosomes 1 and 4. Two mixed-linear model association analyses are shown, overlayed. In the first, GWAS was conducted on IDM (purple) and non-IDM (gray) SNP markers. For the second, GWAS 250 Kb window-based mean *M. glaziovii* allele dosages at IDMs (*DoseGlaz*; orange squares). The horizontal lines represent the bonferroni-significance threshold for the *DoseGlaz* (orange line) and SNP GWAS (purple line). In the bottom left quadrant is a boxplot of all bonferroni-significant marker-effects pooled by trait and chromosome

On chr. 1 between 24.0-31.9 Mb significant IDM and *DoseGlaz* were detected for DM (mean effect of *M. glaziovii* alleles in percent DM: 1.05 IDM, 1.49 *DoseGlaz*) and RTNO (mean effect in ln(kilograms/plot): 0.08 IDM, 0.09 *DoseGlaz*) (Table S18). For MCBSDS the Chr. 4 QTL includes *DoseGlaz* and IDM, covering most of the introgression region, from 12.6-23.4 Mb. For SHTWT and HI however, the region spanned only from 22.35-25.1 Mb and there was a single significant marker for RTNO and RTWT nearby at 17.9 Mb. Effects on chr. 4 of *M. glaziovii* alleles for brown streak disease appear protective (mean effect on disease severity [1-5] score: −0.17 MCBSDS, −0.33 CBSDRS). For SHTWT (units: ln(kg/plot)) and HI (units: proportion 0-1) mean *M. glaziovii* effects were −0.085 and 0.023 respectively. In addition there was one *DoseGlaz* significant for MCBSDS on chr. 5 and one on chr. 12. The sig. *DoseGlaz* on chr. 12 was estimated to *increase* disease susceptibility with an effect-size of 1.22 (trait scale 1-5). Note that RTNO and SHTWT effects are on the natural log scale.

### Impact of introgressions on genomic prediction

Genomic selection (GS) is becoming an important part of modern cassava breeding (16, 38). We investigated the impact of introgression regions on genomic prediction accuracy, which is directly proportional to their contribution to breeding gains during GS, all other things being equal. We did ten replications of five-fold cross-validation for each trait-institute dataset. We tested five prediction models: non-partitioned (ALL markers), genome-partitioned and IDM-null models. For the “genome-partitioned” and “IDM-null” models, we divided markers into two kinship matrices either randomly or based on whether a SNP was an IDM or not.

The accuracy of partitioned models was almost identical to the non-partitioned model for both the IDM-based and the random genome-partitions. However, removing the IDM-based component from the model tended to reduce accuracy, especially in the NaCRRI data, on average 0.004 (max 0.04) relative to the PARTITIONED and 0.005 (max 0.03) relative to the ALL models (Fig. 6A). These comparisons provide a means to measure the importance of introgression regions in ongoing GS. In contrast to the IDM-based genome partition, removing the equivalent random components decreased accuracy an average 0.001 (compared to ALL) and −0.001 (compared to PARTITIONED) (Fig. S15, Tables S19-20).

**Fig. 6.**
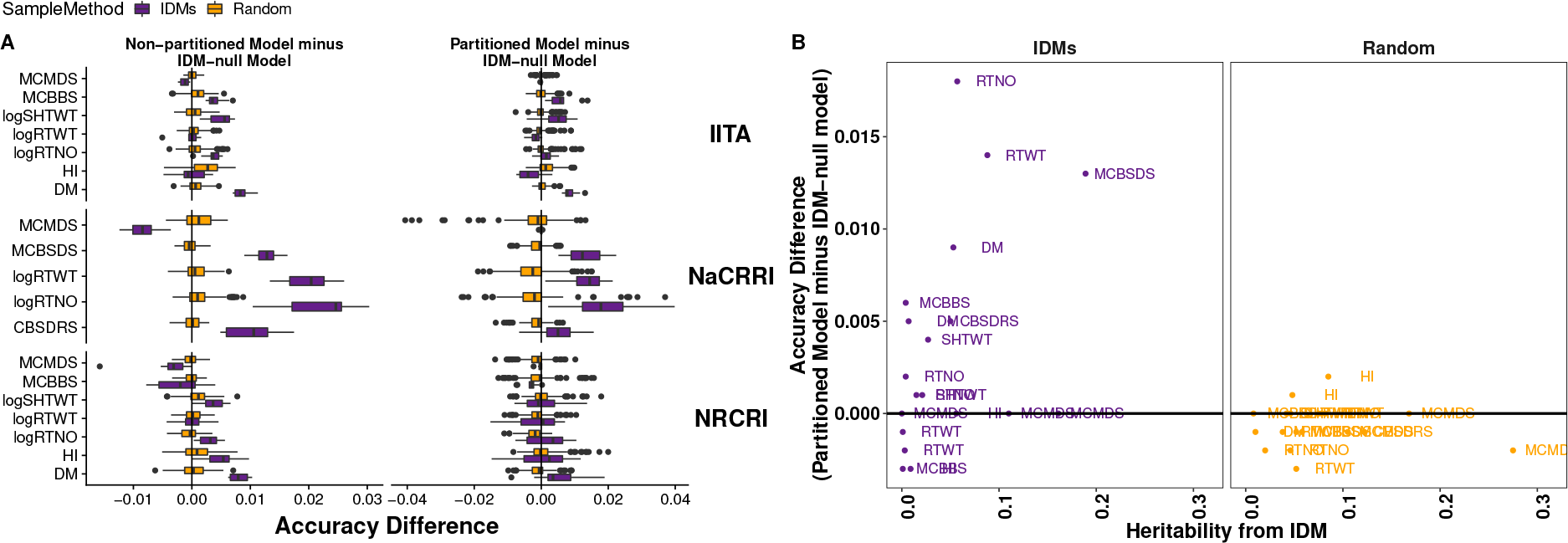
The importance of introgressions for genomic prediction. (A) The difference in prediction accuracy between a model with vs. without the introgression regions is expressed in horizontal boxplots. Ten replications of five-fold cross-validation was conducted for each Trait-Institute combination. For each trait-institute dataset, we used the same 10 random partitions of training-test for each model tested. Two measures are shown on the x-axis: the total accuracy of the partitioned model minus the IDM-null model (A, left), the accuracy of the non-partitioned model minus the IDM null model (A, right). Two methods of partitioning the genome were compared: the IDM-based partition (purple boxplots), and 15 different random partitions, pooled in the (orange) boxplots. (B) The mean difference in prediction accuracy between the partitioned model and the IDM-null is plotted (y-axis) against the introgression-associated heritability (x-axis) from the multi-trial analyses. Results are shown for the IDM-based partition of the genome (purple, left panel) and three tested random partitions (orange, right panel).

Finally, we observed that the size of the impact on prediction accuracy (measured from comparing ALL and IDMnull models) scaled with the 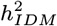 with a correlation of 0.41 for the IDM-based genome partition and −0.09 for the random partition (Fig. 6B).

## Discussion

### Beneficial effects of introgressed alleles are consistent with divergence in their frequencies across the African continent

The original impetus for interspecific hybridization at Amani (circa 1930’s) was to combat cassava mosaic disease (17, 18, 20, 22, 23). We observed consistent and beneficial *M. glaziovii* allelic effects, however, we found neither a beneficial effect nor a significant genetic variance for CMD. In a previous article, focused on GWAS for CMD, we noted an absence of major effect QTL other than CMD2, a dominant, possibly multi-allelic locus (39). We verify here that the protection derived from the CMD2 locus did not arise from introgression as the only associated GWAS result on chr. 12 indicated the *M. glaziovii*-allele increased susceptibility (Fig. 5; Table S18). Introgression-derived CMD resistance has previously been suggested to be weak (relative to CMD2), “recessive” and “polygenic” (25, 26); our results seem to be in agreement with this assessment.

Introgression alleles we did detect at QTL are adaptive and consistent with the population structure we observed (Fig. 1A), arising primarily due to segregation of the two very large segments detected on chr. 1 from 25Mb to the end and chr. 4 from 5Mb to 25Mb, as well as a segment on chr. 14 (Fig. 1B, Fig. 2,3). *M. glaziovii* segments are common in African breeding germplasm (Fig. 2), less common among African landraces and nearly absent from Latin American cassava. Dry matter alleles from *M. glaziovii* at a previously identified QTL on chr. 1 (Fig. 5; (32)) seem to explain the higher frequency of those introgression segments in W. Africa, given the breeding emphasis placed there on that trait as well as yield. The chr. 4 segment, in contrast, is more common in east Africa, which also aligns with the focus there on CBSD resistance breeding (27, 40) and the protective *M. glaziovii* alleles there for that disease (Fig. 5; (30, 31)). We note that, in line with a recent study of cross-continent prediction of CBSD resistance (41), the existence in West Africa of the potentially-protective chr. 4 segment is promising for the possibility to preventatively breed for CBSD resistance in W. Africa. This leads to the reciprocal suggestion that any beneficial DM alleles being targeted in W. Africa are likely to be present and thus potentially useful in E. Africa.

By comparison of African to Latin America clones, we believe our evidence supports the origin of the chr. 1 and 4 *M. glaziovii* introgressions African, in line with historical and recent genomic evidence (29). We do note five CIAT clones with signatures of introgression; one is BRA534, which at 34% *M. glaziovii* genome-wide likely has recent (non African) wild ancestors, and four others were heterozygous for the same segments on chrs. 1 and 4 that the African germplasm have. To date, we have not been able to trace the pedigree or otherwise ascertain the origin of these clones.

### Inbreeding depression due to linkage drag accumulating genetic load in introgression regions may explain homozygote deficit among landraces and elite cultivars

The suppression of recombination, often due to structural variants like inversions, is often a consequence of hybridization between crops and their wild relatives (10, 12, 42). Though we do not know whether an inversion or other structural variant underlies *M. glaziovii* introgressions in cassava, we estimated that recombination is clearly reduced in the introgressed regions on chr. 1 and 4 by 14% and 71%, respectively compared to the rest of the genome ((29, 32); Figs. S6 & S9, Table S21). Further, adjusting for LD using LDAK almost uniformly reduced the heritability accounted for by introgressions (Figs. S12-13). Also, though the introgressions were clearly important for genomic prediction, their overall effect on accuracy was small (Fig. 6; Fig. S15). This suggests that while introgressions are clearly still important, having uniformly beneficial effects at QTL (Fig. 5) and nearly doubling in frequency during three cycles of GS (Fig. 3), their physical size is disproportionate to their true economic value.

One theoretical prediction about introgressed alleles under selection with suppressed recombination is that they can result in the accumulation genetic load due to linkage drag (10, 11). This is especially a concern for vegetatively-propagated non-inbred crops like cassava (33). We observed that putatively deleterious alleles in introgression regions accumulated relatively faster (both LG to GG and GG to C1) compared to the rest of the genome. We further observed balancing selection in the form of an *M. glaziovii* homozygote deficit from variety trials.

In clonally-propagated crops, selection for advancement during variety trials is using total genetic merit rather than breeding value based on performance in a series of field trials with progressively more replicates, locations and increasing plot size. The GS progeny that we analyzed (and thus the sample in which we observed an initial increase in *M. glaziovii* homozygosity) represent those that successfully germinated and were vigorous enough early on to warrant genotyping. We discovered that *M. glaziovii* homozygotes were excluded from later stage field trials early in the process (Fig. 3D). This indicates there may be phenotypically-expressed negative effects of these introgressions, which may be related to the accumulation of homozygous deleterious mutations we observed. Linkage drag in adaptive introgression regions has been proposed to explain balancing polymorphism regions in cases including human-Neanderthal hybridization (2) and wing mimicry in *Heliconius* butterflies (43, 44).

Introgression-associated inbreeding depression is thus a critical topic for future investigation. At present, cassava breeders are maintaining introgression heterozygosity at great cost, through a multi-stage selection process. First, favored crosses between heterozygous parents generates many unsuitable offspring, homozygous for introgressions and suffering some as yet unquantified inbreeding depression. Subsequently, field evaluations are required to identify and purge these individuals. We propose that targeted recombination of introgression segments would increase the rate of gain and sustainability of cassava breeding by allowing simultaneous fixation of beneficial alleles and purging of genetic load.

Taken together, our results point to the continued importance of wild alleles in cassava, one of the most important staple foods in the developing world, and a model for other clonally-propagated root and tuber crops. We present an example of both the benefits and consequences of historical introgression for modern crop breeding. Our methods and the breeding implications we highlight will therefore provide a valuable example for other crops.

## Materials and Methods

Detailed materials and methods are described in the online **SI Appendix**. Raw data and analytic results as well as high resolution maps of introgressions are available here: ftp://ftp.cassavabase.org/manuscripts/.

## ACKNOWLEDGMENTS

We give special thanks to Simon E. Prochnik & Jessen V. Bredeson for conceptual support during initial development of the study. Thanks to Lukas Mueller and Prasad Peteti for data hosting and curation respectively. Thanks also to Luis Augusto Becerra Lopez Lavalle and the International Center for Tropical Agriculture (CIAT) for contributing their germplasm collection for genotyping. Finally, we are grateful to entire Next Generation Cassava Breeding team (www.nextgencassava.org), so many of whom have contributed to this study in the field, in the lab and beyond. We acknowledge the Bill & Melinda Gates Foundation and UKaid (Grant 1048542) and support from the CGIAR Research Program on Roots, Tubers and Bananas.

