## Supplementary material for "Introgressed *Manihot glaziovii* Alleles in Modern Cassava Germplasm Benefit Important Traits and Are Under Balancing Selection": SI_Appendix

#### This PDF file includes:

Figs. S1 to S15
Captions for Databases S1 to S21
References for SI reference citations

#### Other supplementary materials for this manuscript include the following:

Databases S1 to S21

### Materials & Methods

Raw data and analytic results as well as high resolution maps of introgressions are available here: ftp://ftp.cassavabase.org/manuscripts/.

**GBS and WGS Datasets.** In this study, we focused on detecting *M. glaziovii* genome-segments introgressed into cultivated cassava (*M. esculenta*) germplasm and estimating their phenotypic effects.

The HapMapII dataset is a collection of 30X whole-genome sequences (WGS), which has been previously described (1). In our study, we use HapMapII as the basis for identifying introgression diagnostic markers (IDMs), which are described in detail below. The rest of the germplasm we analyzed were genotyped using the genotyping-by-sequencing (GBS) approach (2, 3). The overall GBS protocol and bioinformatics pipeline we employed for quality control and genotype imputation have been described previously (3–5). The specific details of the data are described hereafter.

We have included GBS genotypes and phenotypes (described below) from three cassava breeding programs: National Root Crops Research Institute (NRCRI, Umudike, Nigeria), International Institute of Tropical Agriculture (IITA, Ibadan, Nigeria), National Crops Resources Research Institute (NaCRRI, Namulonge, Uganda).

The genomic selection (GS) training populations for the three programs, IITA, NRCRI (NR) and NaCRRI (UG), have all been previously described (4–12). Additional datasets were sourced from IITA's Genetic Gain (GG) (13) and Local Germplasm (LG) populations, which are also compared herein. The LG is a collection of landraces and local varieties collected mostly, but not exclusively, in west Africa. IITA also contributed GBS data from a panel of Latin American accessions obtained through its collaboration with the International Center for Tropical Agriculture (CIAT). Herein, this dataset is used to compare introgression between African and American germplasm. Finally, in order to investigate change in introgression frequency due to (genomic) selection, we included GBS data from three consecutive IITA progeny generations (hereon C1, C2 and C3), with C1 being descended from selected GG parents, C2 being descended from crosses among selected C1 parents and so on (4).

With the sole exception of the CIAT collection, our GBS samples were derived from the following bioinformatics pipeline: A reference panel of 4629 accessions, consisting of all African germplasm available that were not classified as genomic selection progenies (i.e. three training populations plus assorted landraces) was assembled. Sites were removed if $> 80\%$ had zero reads. Imputation of the "reference" panel was done with Beagle v4.0. The imputed dataset was then filtered post-imputation, keeping sites only if they had allelic R-square (INFO/AR2 field of the VCF) $\geq 0.3$. The imputed and filtered reference set was then used to impute the remaining IITA GS progeny.

The CIAT lines were part of separate study and thus they were processed and imputed along with additional samples (not published). For that dataset, pre-imputation genotype calls were allowed only if a minimum of one read and a maximum of 50 reads were present for each individual at each site. We removed loci with greater than 80% missing and an average mean depth greater than 120. We also thinned markers within 5 bp of each other. These quality control procedures were implemented using the VCFtools (v0.1.14) software package (14). We then imputed missing data using Beagle (v4.1) using the *gl* mode, $window = 2250$, $overlap = 225$ and $niterations = 10$ (15, 16). Post-imputation, we removed markers based on the allelic $R^2$ ($AR2 < 0.3$) statistic (17).

**Introgression Diagnostic Markers.** We used an ancestry informative marker approach, similar to that of (18) in order to detect introgressed *M. glaziovii* genome segments. Our approach relies on the comparison of a set of pure (i.e., non-introgressed) *M. esculenta* (Me reference) to a collection of *M. glaziovii* (Mg reference). From the comparison of these two reference panels, we identified a set of introgression diagnostic markers (IDM) that can be used for detecting *M. glaziovii* segments in an admixed sample. We used two criteria to classify a SNP as introgression diagnostic. Either the SNP must be fixed for different alleles between the Me and Mg reference panels ("Strict" IDMs) or the SNP must be fixed in the Me reference but polymorphic in the Mg sample ("GlazPoly" IDMs). The rationale for "GlazPoly" IDMs is that, if we identify alleles that are only present in the *M. glaziovii* panel and not in the pure *M. esculenta*, then finding the *M. glaziovii* alleles at those sites in an introgressed individual would be diagnostic, or at least contribute to our confidence that an *M. glaziovii* genome segment was present.

In order to detect introgression segments in our admixed GBS dataset, we first identified a set of IDMs by analyzing the cassava HapMapII dataset (1). HapMapII contains 238  30X whole genome sequences ( 28M SNPs) including 8 *M. glaziovii*, 11 *M. esculenta* x *M. glaziovii* hybrids, 16 *M. flabellifolia* (wild progenitor of cassava), a few other wild relatives and 200 cultivated *M. esculenta* samples.

The first step towards identifying a set of IDMs was to define panels of *M. glaziovii* and *M. esculenta*. HapMapII includes 16 samples of the wild progenitor of cassava, *M. flabellifolia* We exclude these samples from our study. For our *M. glaziovii* reference panel, we selected seven samples (GLA59008, GLA59008-1, MAN00401, MGLAZIOVII, Mglaziovii, MglazioviiR, MglazioviiS) that were marked as *M. glaziovii* in HapMapII (Supplementary Table 1 of (1) and included one additional sample (IRWA02712), because in admixture and clustering analyses (not shown) IRWA02712, though it was marked as *M. irwinii*, is not distinguishable from *M. glaziovii*.

Defining a reference panel of "pure" *M. esculenta* required greater care than for *M. glaziovii* since we know our sample potentially includes admixed individuals but don't yet know which. In (18), an analysis using the software frappe (19) was the primary basis for defining "pure" cassava. In order to ensure that on a segment-to-segment basis our cassava reference set truly did not contain *M. glaziovii* segments, we defined Me reference panels on a per-window basis. For every 1000 SNPs, we calculated the pairwise Hamming distance (as implemented by the –distance flag in plink1.9beta3.3, www.cog-genomics.org/plink/1.9/, (20)) between all HapMapII samples. In order to make this procedure more computationally tractable, we first LD pruned the 28M

SNP set using the plink1.9 –indep-pairwise flag with a window size of 50 SNPs, step size of 10 and an LD r2 threshold of 0.3. After LD-prunning, the HapMapII dataset had 4,952,655 SNPs left ( 4.95M). For each 1000 SNP window, we calculated the mean Hamming distance between the Mg reference panel and each cultivated cassava sample (MeanGlazDist for short).

Before making decisions about which cassava clones were genetically distant enough from *M. glaziovii* to use as a window-specific reference panel, we plotted summary statistics (Fig. S1). Specifically, we calculated the mean, median, maximum and standard deviation of MeanGlazDist variable for each window (Fig. S2).

We noted some outlier windows at the end of chromosomes where all summary statistics are very close to zero. We exclude these windows from further consider by filtering cases with MeanGlazDist<200. We then select as Me reference for each window, the 10 clones with greatest MeanGlazDist in that window.

In order to apply IDMs for detecting introgressions in the GBS data we restrict our downstream analyses to the 149,098 GBS sites intersecting both GBS and WGS datasets. We conducted a principal components analysis on three marker sets: all markers [IDM+non-IDM], IDM markers and non-IDM markers. Note that for this analysis, non-IDM markers which were later designated as tag-IDM because they are in strong LD with the set of IDM, were only included in the "all markers" analysis. We describe the definition of tag-IDM in detail below under Introgression Tagging Variants. Only the populations UG, GG, NR and LG (N=2742) were included in the PCAs. Only SNPs with minor allele frequency (MAF) > 0.01 were included in the analysis. We used the *prcomp()* R function, with center and scale arguments set to TRUE.

The PCA and many other downstream analyses were conducted on allelic dosage matrices, where the genotypes for each individual (rows) and each SNP (columns) is represented as 0, 1, 2. Note that our dataset contains non-integer values between 0 and 2, representing the uncertainty of imputation. For the non-IDM and tag-IDM, the counted allele is the default for the *plink "–recode"* function. We coded the IDM dosages so the counted allele is the *M. glaziovii* diagnostic allele (*plink* "–recode-allele"). Doing this enables us to interpret eigenvector coefficients (PC loadings) *and* SNP effects (see re: GWAS below) for those markers relative to introgressions. For example, a positive loading for an IDM SNP means individuals at the high end of that principal component are more likely to have an *M. glaziovii* diagnostic allele at that site than individuals at the low end.

**Mapping introgressions in windows.** GBS (2, 3) produces genotype data with a high proportion of missing sites and a low average read depth, which necessitates imputation (15, 21) for most applications (see above).

Individual IDM genotype calls may be incorrect. This means the use of IDM to detect introgressions have some probability of both false positives and negatives. One step we took to reduce the potential noise from individual IDMs was to follow a window-based approach similar to Bredeson et al. (18). We computed the average *M. glaziovii* dosage across IDMs in 250 Kb , non-overlapping windows across the genome (referred to also as *DoseGlaz*).

The window-based dosages were used to generate genome-wide maps of introgression status for each sample and for GWAS (see below).

**Interpolating marker genetic distances.** Marker genetic distances were interpolated using the consensus genetic map from International Cassava Genetic Map Consortium (ICGMC) (22) based on the cassava reference genome v4 (23). The 22404 ICGMC Markers flanking sequences (105 bp) were filtered on biallelic variation and subjected to a nucleotide BLASTN procedure (24) against cassava reference genome version 6 (18) to obtain their new physical positions. Using a custom python script and these inferred ICGMC v6 positions, we interpolated all GBS markers in our dataset.

**Comparing introgressions among populations.** There were eight datasets (GG, LG, NR, UG, C1, C2, C3 and CIAT) for which we wanted to compare introgression status. We computed both the *M. glaziovii* allele frequency and homozygosity rate. For each population, these summaries were made both on a per IDM and a per individual basis. Further, we examined differences between populations on a genome-wide basis as well as in focal introgression regions on chr. 1 (from 25 Mb+) and chr. 4 (5-25 Mb).

**HAPMIX in HapMapII.** In order to provide additional confidence in the introgressions we detect using IDMs, we applied an alternative approach to the HapMapII dataset. We used HAPMIX (25), a haplotype-based method for local ancestry inference in populations formed by two-way admixture. HAPMIX infers the ancestry of chromosomal segments by extending a population genetics model developed by Li and Stephens (26). The method assumes that the admixed population of interest arose from a single admixture event between two ancestral populations. In our case, the two ancestral populations are *M. glaziovii* and *M. esculenta*. For each chromosome, the method requires (1) phased data from two unadmixed reference populations that are closely related to the true ancestral populations, (2) unphased data from a sample of admixed individuals, and (3) a recombination rate file, listing the physical position and genetic position of each site. HAPMIX first estimates the likelihood that an allele at a given site on an admixed haplotype originates from each ancestral population then uses a Hidden Markov Model (HMM) to combine likelihoods at each site with information from neighboring sites to provide a probability of ancestry at each site. Before running HAPMIX, we used HapCUT (27) on already Beagle-imputed and phased HapMapII dataset in order to improve local phasing.

***Defining the two reference populations and the admixed sample.*** For local admixture analysis, we needed two reference populations: reference population 1, consisting of individuals related to the true *M. glaziovii* ancestral population, and reference population 2, consisting of individuals related to the true *M. esculenta* ancestral population. We used the same eight *M. glaziovii* individuals that were used to define IDMs. To define reference population 2, we selected the 10 *M. esculenta* individuals in HapMapII (out of the 217 *M. esculenta*) that were most genetically distant from the eight *M. glaziovii* samples based on IDMs analysis. They were: CW45617, CPCR15B91, CPCR27B7, CPCR27B17, MBRA685, UG08S0P003, CR4442, CPCR24B3, BRA8565, UG08S0P005. Note that this was in contrast to the window-specific approach to defining *M. esculenta* reference individuals that was used to define IDMs. To define the genome-wide reference, we simply summed the *MeanGlazDist* values across each individual's genome and selected the top ten.

The remaining 207 *M. esculenta* served as our admixed sample. Discrepancies between the reference populations and the true ancestral populations should, in theory, introduce error into the inference procedure; however, HAPMIX has shown to be robust to errors in defining the reference populations as HAPMIX takes into account the possibility of incomplete sampling of diversity and that the reference populations have drifted from the true ancestral populations (25).

***Specification of HAPMIX parameters.*** HAPMIX requires specification of nine model parameters: (1) the average number of generations since admixture, T, (2-3) the rates at which there is copying of ancestry segments from the 'wrong' population, $p_1$ and $p_2$, (4-6) 'mutation' parameters $\theta_1$, $\theta_2$, and $\theta_3$, (7) the probability that a given segment of an admixed haplotype originates from ancestry population 1 and population 2, $\mu_1$ and $\mu_2$, and (8-9) the 'recombination' rate between haplotypes within reference population 1 and 2, $\rho_1$ and $\rho_2$. We selected parameter values via a process of trial and error, running HAPMIX on the 11 *M. esculenta* x *M. glaziovii* hybrids in HapMapII (results not shown). For each hybrid, when tuned properly, HAPMIX should infer the presence of one *M. glaziovii* allele at each site (or at least, a large proportion of sites). After parameter tuning, we selected the following parameter settings for HAPMIX: T = 4, $p_1 = 0.05$, $p_2 = 0.05$, $\theta_1 = 0.2$, $\theta_2 = 0.2$, $\theta_3 = 0.01$, $\mu_1 = 0.2$ (and $\mu_2 = 1 - \mu_1$), $\rho_1 = 700$, and $\rho_2 = 900$. HAPMIX has multiple output format options, specified using the parameters "HAPMIX_MODE", "OUTPUT_DETAILS", and "THRESHOLD". We run HAPMIX using the HAPMIX_MODE = "DIPLOID", OUTPUT_DETAIL = "HAPLOID_FILES", and THRESHOLD = 0.9. These settings tell HAPMIX to output two files per haplotype (i.e., four files per individual): a file with each site's ancestry status (HAPMIX calls a site's ancestry status only if it passes a probability threshold of 0.9), and a file with each site's allele.

### Introgression Tagging Variants.

**Introgression Tagging Variants.** Above, we identified IDMs, then called and quantified introgression segments in the extant germplasm. Our next major objective was to quantify the phenotyping impact of segregating *M. glaziovii* genome segments on the germplasm. One approach uses genomic mixed models to estimate the amount of genetic variance attributable to introgression regions relative to the rest-of-the-genome (28–30). To do this, we first needed to ensure that we were able to obtain independent estimates of genetic variance for introgression vs. non-introgression regions. If unaddressed, LD between IDM and non-IDM SNP sets will lead to non-independent estimates of genetic variance when fitting the model described above (30–33). The variance arising from introgression regions might then be captured by the non-introgression regions and vice versa.

It is impossible to entirely eliminate this problem: long distance LD exists in populations of clones driven by population structure and familial relatedness. However, to reduce the non-independence, for every SNP not previously identified as IDM, we calculated two statistics. Both are based on pairwise LD $r^2_{LD}$ statistics as computed in R as the squared correlation among allelic dosages. From these $r^2_{LD}$'s, we first determined the maximum LD (maxLD) observed between each non-IDM SNP and the set of IDMs. Our rationale here was that non-IDM SNPs in *very high* LD with even one IDM SNP could explain genetic variance attributable to the same causal variants. Second, we calculated the total LD (totalLD) between each non-IDM and the entire set of IDMs. This metric is essentially the same as the LDscore, which is calculated on a window-basis (30–33). The totalLD, we reasoned, might provide an even better idea (compared to maxLD) of the degree to which an non-IDM SNP "tagged" the introgressed regions of the genome.

We needed an at least semi-objective approach to choose a threshold for declaring SNPs as "tagging" IDMs or not. We tested a range of LDscore (300-1500, interval 200) and maxLD (0.1-1, interval 0.2). For each threshold, we partitioned the SNPs into IDM, non-IDM and tag-IDM accordingly. We then constructed 4 kinship matrices using *A.mat* function from the rrBLUP R package (34): tag-IDM, IDM, non-IDM and IDM+tag-IDM. We used the correlations between the upper off-diagonals as a proxy for the independence of genetic variance components that might result. Our objective was therefore to partition the SNPs using LDscore and/or maxLD such that we maximize the cor(tag-IDM, IDM) and minimize both cor(tag-IDM, non-IDM) and cor(IDM+tag-IDM, non-IDM). The correlation between the kinships using the original partition of IDM and non-IDM (0.37) was the baseline to improve upon.

We chose to use a LDscore threshold of 500, because they were more similar in the kinship they measured to the IDM than to the non-IDM (Figs. 5 -7; Tables S1 & S4). By redesignating these originally non-IDM SNP as tag-IDM and including them in the kinship matrix with IDMs we reduced the correlation of IDM and non-IDM kinships to 0.30. We included tag-IDM in the IDM kinship matrices used in all subsequent analyses. With this procedure, we hoped to improve our ability to distinguish introgression-associated (IDM + tag-IDM) from the rest of the genetic variance (non-IDM) in key cassava traits. It revealed many regions where LD between non-IDM and IDM markers was elevated (Fig. S6).

### Field Trials.

***Trials chosen.*** For this study, we compiled data from 68 field trials (42 IITA, 5 NaCRRI, 21 NRCRI), which were scored for nine traits. NaCRRI, NRCRI and IITA (see **Datasets** above) have genomic selection (GS) programs as part of a project called Next Generation Cassava Breeding (www.nextgencassava.org). For the NR and UG populations, the trials included in our analyses comprise the genomic selection training populations (TPs). For IITA, trials from both the GG (original TP) and LG (landrace / local germplasm) populations were included. There were a total of 2742 phenotyped clones in the dataset.

With the exception of the LG dataset, versions of most of these data have been analyzed in other publications (4, 7–9, 12). Versions of these trials are available from the online database www.cassavabase.org. The data used in this study are freely available here: ftp://ftp.cassavabase.org/manuscripts/. In addition, we summarize the trials in terms of the number of observations (Nobs), clones (Nclone), reps (Nrep) and the ratio of Nobs/Nclone (ObsToCloneRatio) per-Institute-per-Trial (Table S6).

***Traits scored.*** Cassava faces a number of pest and disease problems in Africa (35). We included severity scores, which are on a standard scale of 1 (no symptoms) to 5 (very severe symptoms), for three diseases: cassava brown streak disease root necrosis (CBSDRS) and foliar (MCBSDS), season-wide mean cassava mosaic disease (MCMDS) and cassava bacterial blight (MCBBS).

We also scored five yield-related traits: dry matter content (DM), fresh root weight (RTWT), fresh shoot weight (SHTWT), root number per plot (RTNO), and harvest index (HI). MCMDS and MCBSDS are the mean of measurements taken at up to three time points throughout the season: 1, 3 and 6 MAP. Dry matter content is the percentage of dry root weight relative to fresh root weight (RTWT). At IITA, DM was measured by drying 100g of fresh roots in an oven whereas at NRCRI and NaCRRI, the specific gravity method ((36)) was used. Both RTWT and SHTWT were expressed in kilograms per plot. The HI was the ratio of RTWT to RTWT plus SHTWT. RTNO was the number of roots harvested from each plot. For all analyses below, RTNO, RTWT and SHTWT were natural-log transformed to improve homoscedasticity of residuals. Refer to the Cassava Trait Ontology (http://www.cropontology.org/ontology/CO_334) and our previous publications for additional details (e.g. (4, 5). There were as many as 68 trials scored for MCMDS, and as few as 5 for CBSDRS/MCBSDS (NaCRRI only) (Table S6).

**Genetic Variance from Introgressions.** We used a linear mixed-model to model the field trial data described above (37, 38). When the genotypic effect is modeled as random, with an assumed covariance proportional to the coancestry coefficient, which is its associated variance component and is an estimate of the additive genetic variance (and hence the heritability, $h^2$) (28, 30, 39). The genetic covariance matrix can be derived from its expectation based on pedigree (a.k.a. the numerator relationship matrix) or genome-wide markers (a.k.a. genomic realized relationship matrix). We start with the basic mixed model of form described above:

$$y = X\beta + Zg + \epsilon$$

In this model, $y$ is a $n$ (observations) x 1 vector of phenotypic records, $X$ is the $n$ x $p$ design matrix relating observations to corresponding $p$ levels of the fixed-effects. The $p$ x 1 vector $\beta$ contains the fixed-effect estimates. The $n$ x $q$ design matrix $Z$ related the records in $y$ to the $q$ levels of the random effects vector $g$, in this case $g$ is the genotype (unique cassava clone) effect. The $n$ x 1 random vector $\epsilon$ is the residual or error term.

We assume the following about the random effects vector:

$$\begin{pmatrix} g \\ \epsilon \end{pmatrix} \sim N\left(0, \begin{bmatrix} \sigma_g^2 K_g & 0 \\ 0 & \sigma_\epsilon^2 I \end{bmatrix}\right)$$

Both random effects have a mean of 0. The genomic relationship matrix $K_g$ is a square symmetric covariance matrix. It was calculated from genome-wide SNP markers, the same as used for the PCA (see above). We used the first method from (40) as implemented in the *A.mat()* function in the *rrBLUP* R package (34). is the identity matrix, which specifies the usual independent and identically distributed constraint on the residuals. Given $K_g$, $\sigma_g^2$ is an estimate of the additive genetic variance and $g$ (the best linear unbiased predictors, BLUPs) are, in this case, often called genomic-estimated breeding values (GEBVs).

We want to partition the total heritability ($h_g^2$) into a component due to the regions with introgressed *M. glaziovii* alleles ($h_{IDM}^2$) and a component attributed to regions without introgressions ($h_{nonIDM}^2$). We do this by constructing two GRMs, one with markers from IDMs plus tag-IDMs ($K_{IDM}$) and the other with the rest of the markers ($K_{nonIDM}$). We fit the following, expansion on the mixed-model above:

$$y = X\beta + Zg_{IDM} + Zg_{nonIDM} + \epsilon$$

$$\begin{pmatrix} g_{IDM} \\ g_{nonIDM} \\ \epsilon \end{pmatrix} \sim N\left(0, \begin{bmatrix} \sigma_{IDM}^2 K_{IDM} & 0 & 0 \\ 0 & \sigma_{nonIDM}^2 K_{nonIDM} & 0 \\ 0 & 0 & \sigma_\epsilon^2 I \end{bmatrix}\right)$$

Here $Z$ is the design matrix for both $g_{IDM}$ and $g_{nonIDM}$ as long as the order of the rows/columns of the two corresponding kinship matrices is the same.

For simplicity, we will sometimes refer to the two models described above as the ALL and the PARTITIONED models, respectively.

***Per-trial analysis.*** Our first analysis was on a per-trial basis, where a "trial" is defined as a unique experiment planted in a single location-year.

In addition to the two genetic models described above, for each trial, we also fit three additional models:

**IID:** $y = X\beta + Zg_{IID} + \epsilon$, $g_{IID} \sim N(0, \sigma_{IID}^2 I)$

**IDM:** $y = X\beta + Zg_{IDM} + \epsilon$, $g_{IDM} \sim N(0, \sigma^2_{IDM} K_{IDM})$

**IDMnull:** $y = X\beta + Zg_{nonIDM} + \epsilon$, $g_{nonIDM} \sim N(0, \sigma^2_{nonIDM} K_{nonIDM})$

We also fit a NULL, with no genetic component. In some cases, for the NULL model, when no non-genetic effects were relevant, we fit an intercept, or an intercept + NOHAV fixed-effects model, using the *lm* function in R (v3.4.3). For the rest of the models described above, we fit them with the *mmer* mixed-model solver function in the *sommer* (v3.1) R package ((6, 41).

Since a trial is a single location-year, the only fixed-effect was the number of plant stands harvested (NOHAV), fit for RTWT, RTNO and SHTWT (6) . Non-genetic random effects were added, where relevant, on a trial and breeding program-specific basis. Replication effects were fit to replicated trials for all three breeding programs. For NRCRI and NaCRRI, two random effects, one for complete blocks (replication effects) and another for incomplete blocks nested within replications. All non-genetic components were assumed to be *i.i.d.* (covariance equal to $I$).

For model comparisons, described below, we manually calculated the Akaike Information Criterion (AIC) as $AIC = 2 * npar - 2 * logLik$, where $npar$ is the number of fixed+random parameters fitted and $logLik$ is the log-likelihood from *sommer*'s solution.

In the case of nested model comparisons described below, we conducted likelihood ratio tests (LRT) to determine the significance of individual random effects. Using the likelihoods from the models describe above, we compared $2 * (logLik_{full} - logLik_{null})$ to a chi-square distribution with $df = npar_{full} - npar_{null}$. Here the *null* model refers to a model "nested" within the*full* model, i.e. with one or more random-effect dropped.

Each genetic model was compared to the NULL (non-genetic) model ($LRT_{null}$).

***Multi-trial analysis.*** Using the results of the trial-by-trial modelling, we first flagged any trait-trial for which there wasn't at least one genetic model with significant ($p_{LRT_{null}} < 0.05$) and removed them. Next, among the remaining trials, we removed any that didn't have at least one of the non-IID (genomic) models significant (again $p_{LRT_{null}} < 0.05$). The test $LRT_{null}$ only indicates that the two models being compared are different, not which one is actually better. Therefore, among the remaining trials, we used AIC to determine in which case any of the genomic models were at least as good as the IID model. If ($AIC_{IID}$ was 2 units smaller than $AIC_{genomic}$ we considered the non-genomic model to be better fitting than the genomic one, and subsequently removed those trials.

We used the individual results from per-trial analyses, as described above, to identify and filter out data with very low genetic signal (Table S6). Having curated our dataset as described above, we combined data across trials (within institutes) to achieve larger sample sizes and more replications per clone.

For each Trait-Institute data chunk, we fit the genetic models ALL, PARTITIONED and nonIDM described above for per-trial analysis . Since the grouped datasets have multiple locations, years and replications, we added *i.i.d.* random effects for location-year-trial (*LocYrTrial*: trial nested in location-year) and location-year-trial-rep (*LocYrTrialRep*: replication nested in trials). In addition to $LRT_{null}$ as described above, we added a test of the significance of the introgression variance ($LRT_{partition}$) by comparing the partitioned model to the nonIDM-only (IDM-null) model.

***Random partitions.*** There are 38K IDMs. Any partition of the genome with this number of SNPs is likely to explain a significant portion of the genetic variance based on that fact alone. We compared the variance explained by the partition according to IDM status to three random partitions of the same size (Tables S7-8).

***LD-adjusted GRMs.*** Several previous studies have shown that the primary introgression regions on chromosomes 1 and 4 are characterized by strong, relatively long-range LD (9, 18). Excessive (or deficient) tagging of some causal polymorphisms relative to others, for a given trait, is known to bias genetic variance estimates (28, 31). One approach to reduce this bias is to downweight the effect on kinship estimates of SNPs in regions with very high LD, and upweight those in lower LD (31). We used the LDAK version 4.9 to calculate LD-adjustment weights (Table S9) and subsequently to construct LD-adjusted GRMs.

We fit the ALL, PARTITIONED and nonIDM-only models again, this time with the LDAK GRMs in order to observe the impact of LD on the partitioned of variance between IDM and non-IDM.

**GWAS.** We conducted two types of genome-wide association analysis (GWAS) in order to identify quantitative trait loci (QTL) attributable to *M. glaziovii* introgressions.

The first GWAS was on the individual SNP markers (both IDM and non-IDM), excluding those with MAF<5%. We ran the mixed-linear model association (*–mlma*) analysis implemented in *gcta* (v1.25.2) (42). Population structure was accounted by a genetic random effect, $g_{nonIDM} \sim N(0, \sigma^2_{g_{nonIDM}} K_{nonIDM})$. For GWAS with *gcta*, population structure was accounted for by $K_{nonIDM}$, which in this case was constructed using gcta *–make-grm-bin* on non-IDM SNP passing *–maf* 0.01 instead of the rrBLUP::A.mat() version used elsewhere. In the first GWAS, we are able to interpret significant IDM SNPs relative to the *M.g.* diagnostic allele. We further complemented this with a second GWAS, conducted on the introgression-segment dosage matrix (*DoseGlaz*) based on the 15-IDM windows described above. We conducted this GWAS in R, fitting a mixed-model in which each 15-IDM window was sequentially tested as a fixed-effect, and a random effect with the same covariance ($K_{nonIDM}$) as in the *gcta* analysis. From the estimated marker effects $\hat{\beta}$ and their corresponding standard errors $se^2_{\hat{\beta}}$, we calculated a Wald test statistic:

$$WaldStat = \hat{\beta}^2 / se^2_{\hat{\beta}}$$

We obtained a p-value by computing the probability of observed Wald-statistic under the upper tail of a $\chi^2$-distribution with 1 degree of freedom.

As phenotypic responses for both GWAS, we supplied BLUPs for each clone. Using the *lmer* function in the *lme4* R package, we fit an IID genetic model ($g_{IID}$ as described for the per-trial analyses. We analyzed the same curated data and modeled the same design-related random effects (*LocYrTrial* and *LocYrTrialRep*) as in the multi-trial analysis.

**Genomic Prediction.** We measured the importance of introgression regions for breeding value prediction with five-fold cross-validation . We fit the non-partitioned (ALL), PARTITIONED and IDMnull models again using *mmer* in *sommer*. We also fit 15 randomly selected partitions of $n_{non-IDM}$ including the three using in the multi-trial analysis. Because we used a two-stage genomic prediction approach for cross-validation, computation was much faster, making it possible for us to test more random partitions. The first stage is fitting the models described above for use as response data in GWAS. The second stage is the genomic prediction step, but we first de-regressed the BLUPs used for GWAS and weight error variances according to a nonlinear function of the reliability ($r^2$) and the heritability ($h^2$). The procedure is described in multiple previous publications (4, 7, 8) and is based off of (43).

The cross-validation was set-up such that each of the trait-institute dataset were divided up into ten different random partitions of 5 approximately equal parts. For each model on each 5-fold partition of the data, five predictions were made in which four-fifths of the clones' phenotypes were included and one-fifth were left out as a test-set. Prediction accuracy was measured as the correlation among the test-set individuals of their BLUP (as used in GWAS) and their GEBV. For the PARTITIONED model, accuracy was measured for the total GEBV (IDM+non-IDM BLUPs).

**Field plot records of GS progeny.** We downloaded records from www.cassavabase.org of all field plots planted as of January 22nd, 2019 for each of the GS progenies (Table S2). We reserve phenotypic records for these field plots for a future study.

**Deleterious mutations in introgression regions.** We extracted the dosage of deleterious alleles at 9779 of the 22495 putative deleterious mutations identified by(1) from a dataset consisting of the LG, GG and C1 where 5.37 million HapMapII SNPs were imputed. GBS data for the LG, GG and C1 were imputed in a single step using IMPUTE2 (44) with the HapMapII serving as a reference panel. IMPUTE2 parameters were set to values similar to those previously used in (45). Briefly, the number of haplotypes used as a custom reference panel was set to 400, the imputation window was set to 5Mb and the genetic position for each of the HapMapII markers were interpolated from the composite map published by the International Cassava Genetic Map Consortium (ICGMC) (22).

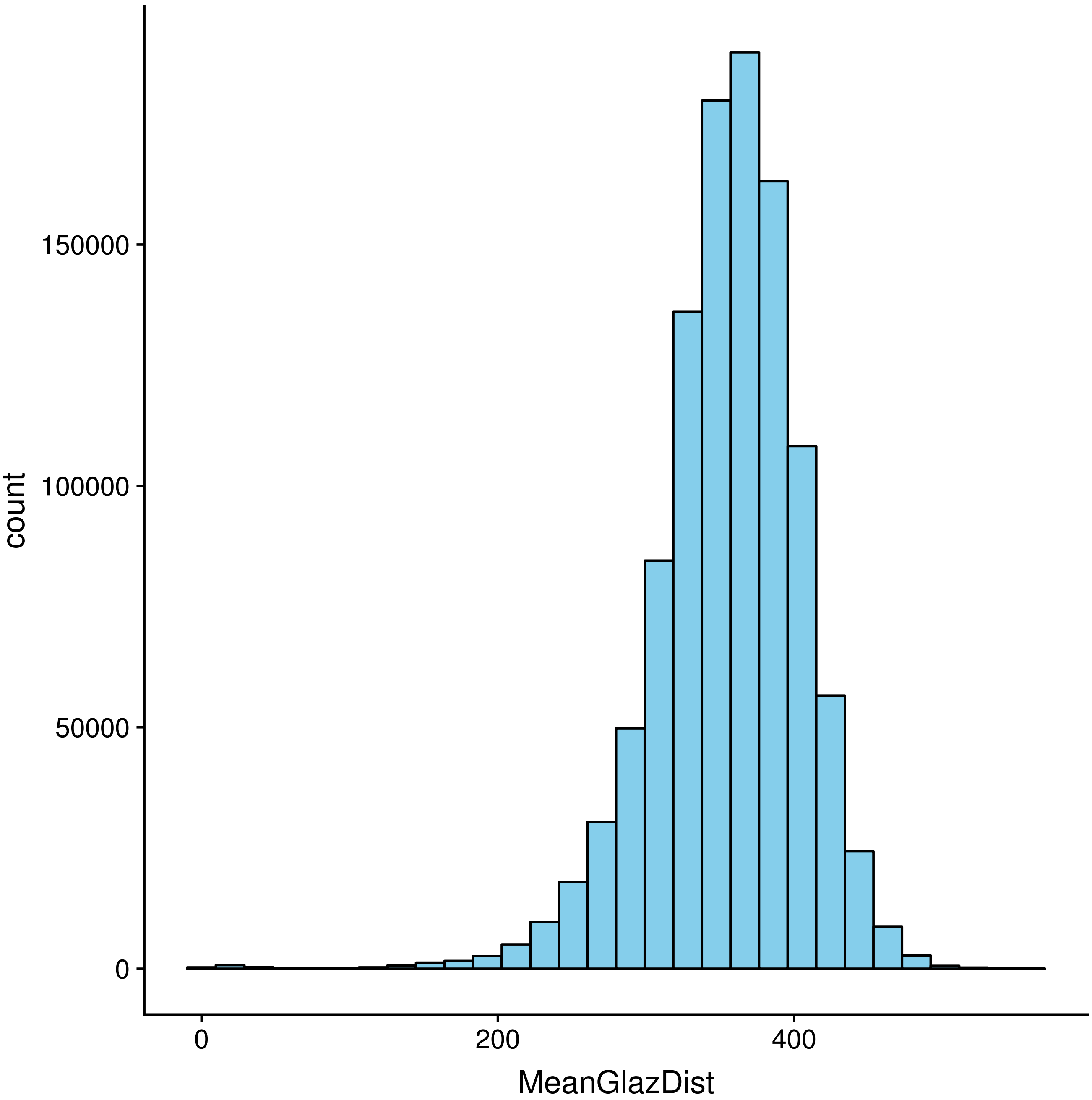


**Fig. S1. Distribution of MeanGlazDist in the cassava portion of HapMapII.** MeanGlazDist = mean Hamming distance for a 1000 SNP window between the *M. glaziovii* reference panel and each cultivated-cassava sample.

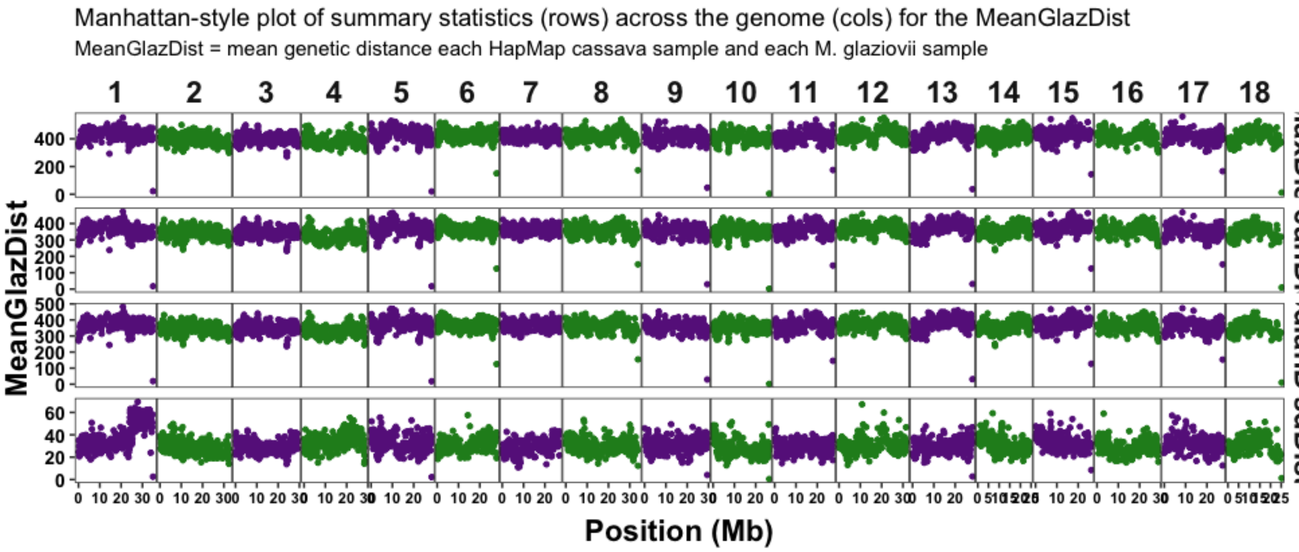


**Fig. S2. Manhattan-style plot of summary statistics (rows) across the genome (cols) for the MeanGlazDist.** MeanGlazDist = mean Hamming distance for a 1000 SNP window between the *M. glaziovii* reference panel and each cultivated-cassava sample.

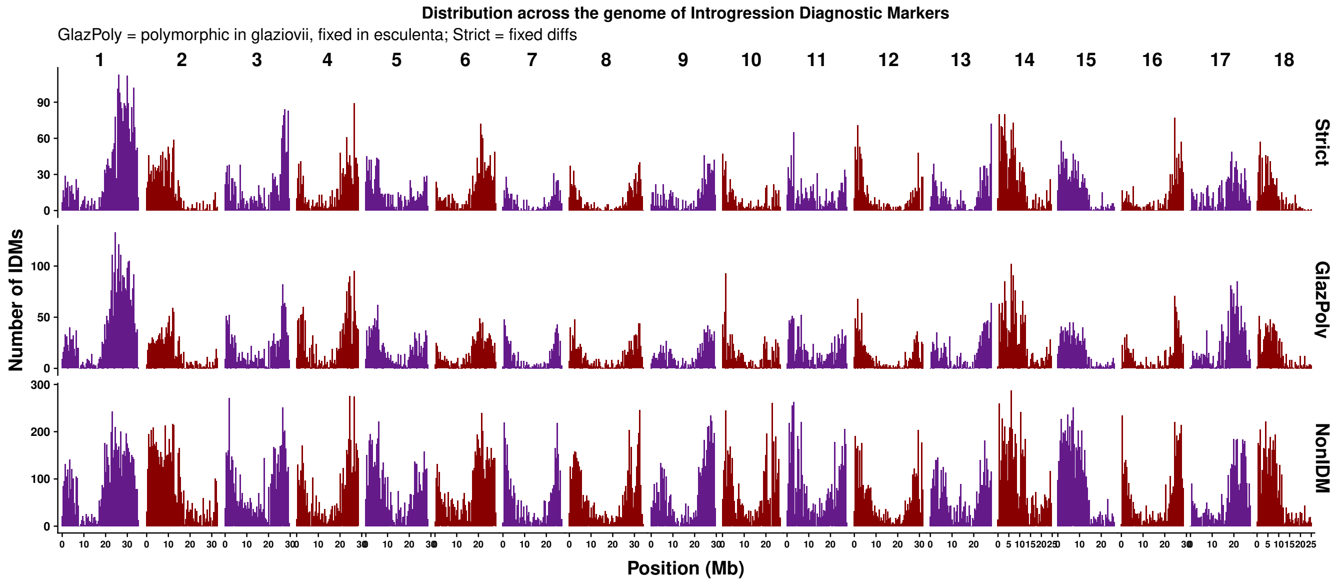


**Fig. S3. Genome-wide distribution of introgression diagnostic markers (IDMs) relative to the rest.** We note from the plot above that the distribution and density of both *Strict* and *GlazPoly*, are very similar to each other *and* also somewhat similar to the *nonIDM* SNPs. The important thing here is that we see that the genome coverage of the diagnostic markers is similar to that of GBS markers in general. We will only be able to reliably call admixed cassava germplasm as introgressed in regions where we have *IDMs*.

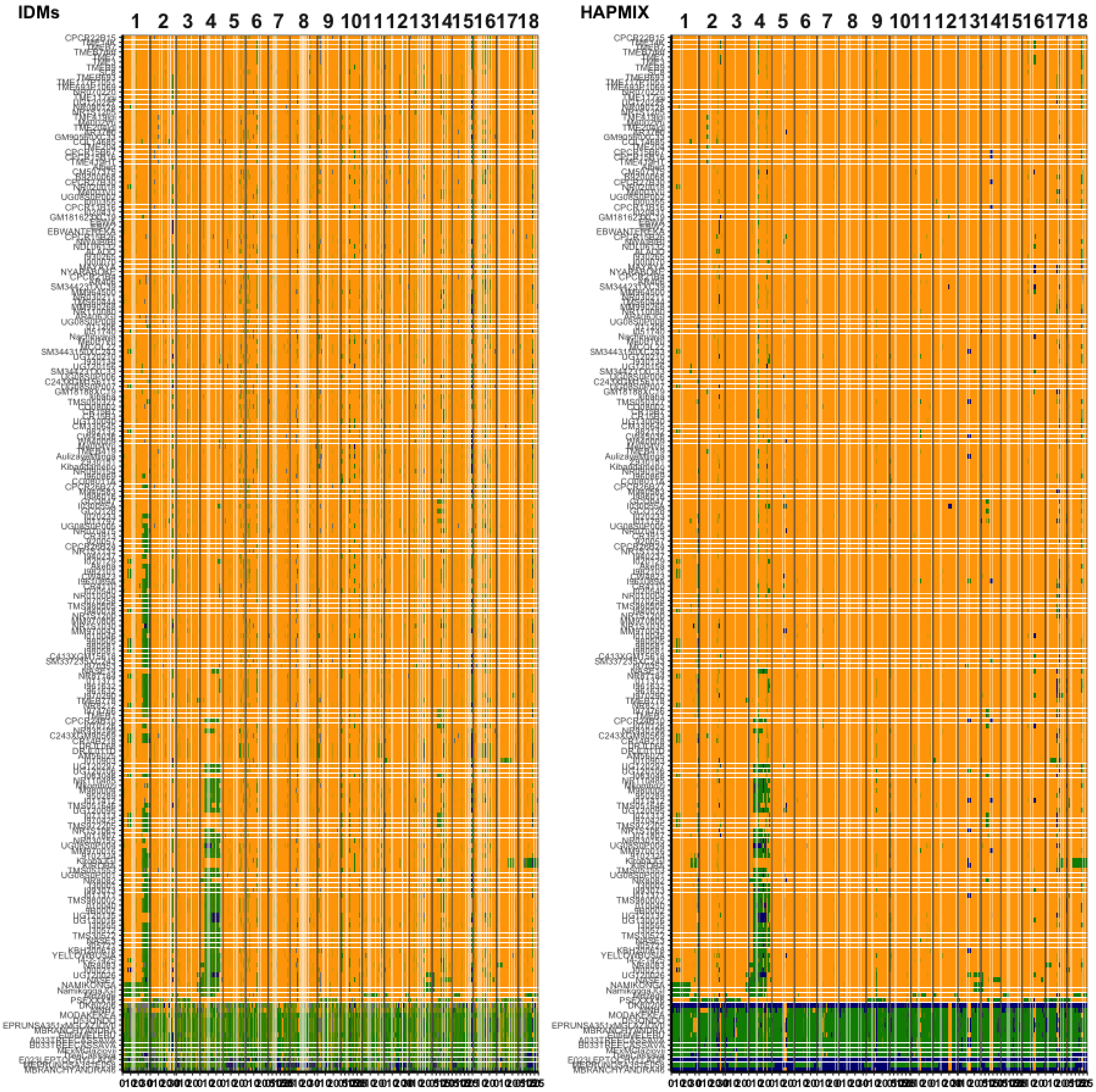


**Fig. S4. Comparison of introgressions detected by IDMs vs. HAPMIX in HapMapII.** The mean *M. glaziovii* allele dosage at IDMs in 250Kb windows across the genome is based on introgression diagnostic markers (IDMs, left) and also based on local admixture modelling (HAPMIX, right). Physical position on each chromosome is depicted in megabases (Mb) along the x-axis. Colors range from orange (0 M.g. alleles), to green (1 M.g. allele), to dark blue (2 M.g. alleles). Rows (clones) are aligned across A and B and sorted within based on the IDM-based genome-wide proportion *M. glaziovii*.

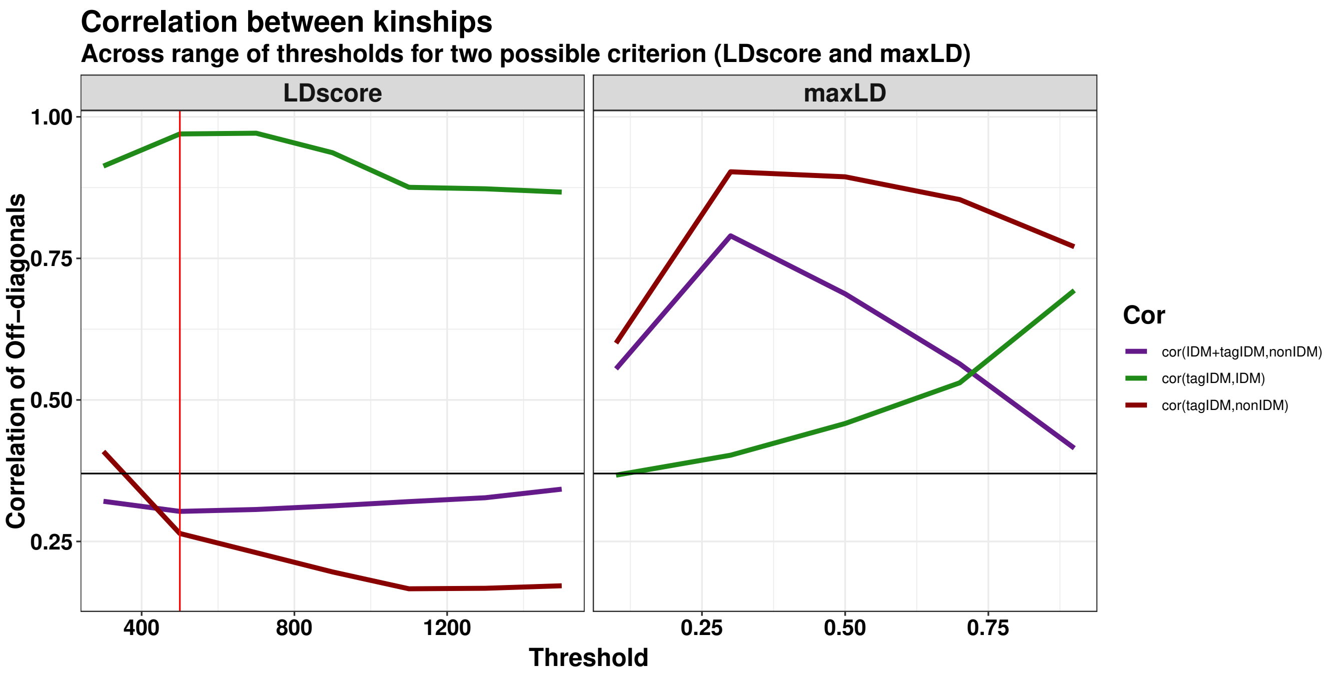


**Fig. S5. Correlations among the off-diagonals of pairs of kinship matrices.** The plot above shows three correlations between the off-diagonals of pairs of kinship matrices created using different sets of SNPs (y-axis). The SNP sets were defined based on a range of thresholds (x-axis) for two criteria for thresholding (panels). The correlations are: cor(IDM+tagIDM,nonIDM), which we want to minimize (purple), cor(tagIDM,IDM) which we want to maximize (green) and cor(tagIDM,nonIDM), which we also want to minimize (red). The horizontal line is the correlation (0.37) between the kinships created for the two original categories (IDM vs. nonIDM), without considering LD. The vertical red line indicates the threshold and the criterion that we ultimately chose to use in order to define addition nonIDM SNPs that "tagged" the introgression regions. The maxLD criterion seems to generally produce correlations between IDM and nonIDM that are greater than the original categories, therefore we rejected that option. In contrast, the LDscore criterion did very well. The cor(IDM+tagIDM,nonIDM) was always lower than the unmodified categories, the cor(tagIDM,IDM) increased to nearly 1 with LDscore thresholds <600 but still remained high across the range and the cor(tagIDM,nonIDM) dropped very low as LDscore-threshold was increased. We chose to use the LDscore criteria with a threshold such that nonIDM SNPs with LDscore>500 were designated as tagIDM and included with the IDM SNPs in downstream analysis (calculation of kinship matrices for variance partitioning).

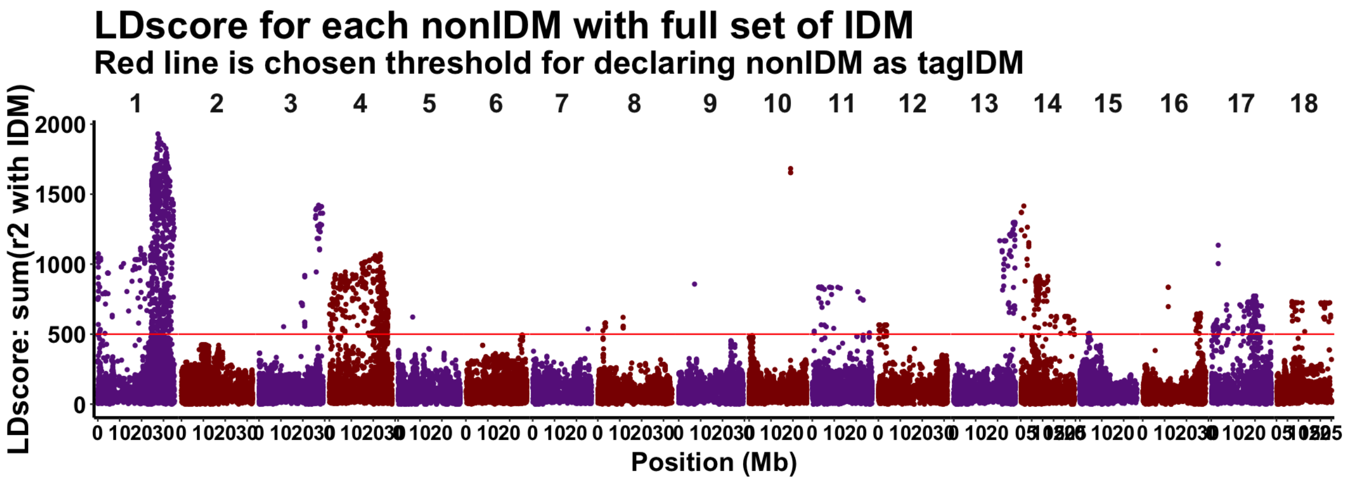


**Fig. S6. LDscore for each non-IDM with the full set of IDM.** The plot above shows the LDscore's (sum of $r^2_{LD}$) of nonIDM SNPs (y-axis) versus their position in the genome (x-axis). The red horizontal line indicates the threshold above which nonIDM SNPs were designated tagIDM.

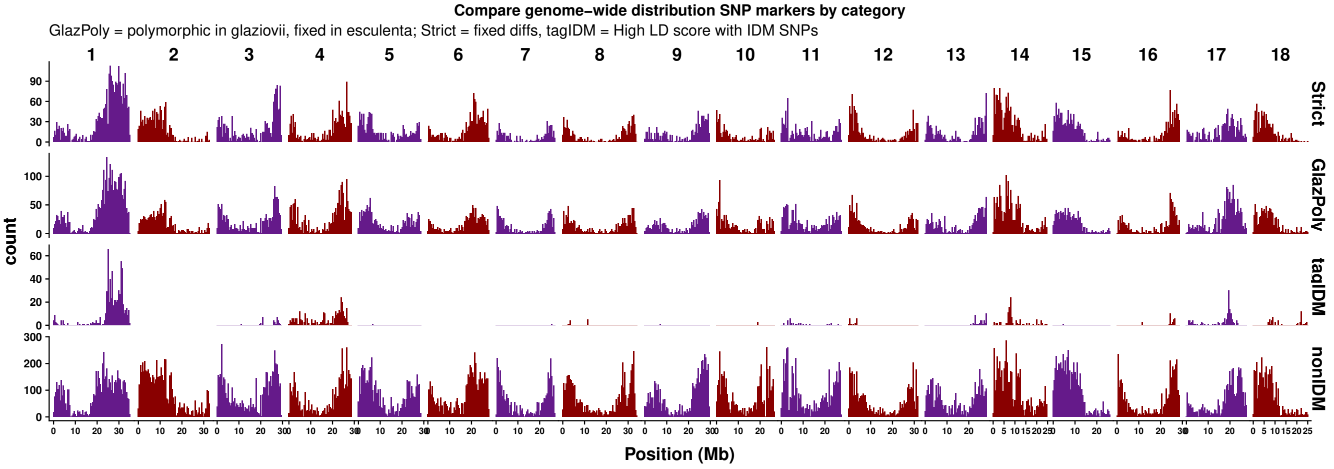


**Fig. S7. Genome-wide distribution of introgression diagnostic markers (IDMs) including tag-IDMs.** Shows the distribution and density along the genome of IDMs both *Strict* and *GlazPoly* as well as *tag-IDMs* and *non-IDM* SNPs.

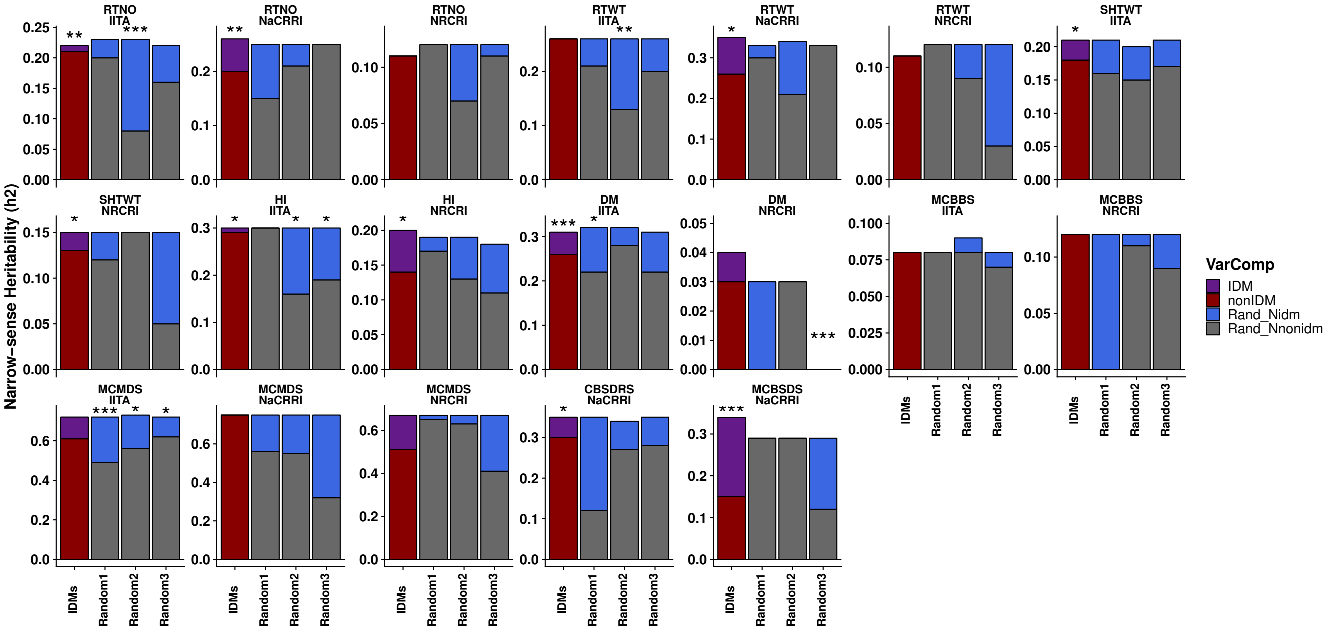


**Fig. S8. Heritability attributable to IDM vs Random genome-partitions.** Each panel shows, for one trait measured at one breeding program, the heritability (y-axis) measured partitioning the genome either based on IDMs or else according to 3 random samples of equivalent number (38000) to the IDMs (x-axis). Heritability was estimated from partitioned genomic mixed-models and the portion of heritability attributable to $N_{IDM}$ component (IDM=purple, Random=royal blue) vs. the rest of the genome (IDMs=dark red, Random=gray) is shown. Stars atop bars represent the level of significance in a likelihood ratio test for the significance of the $N_{IDM}$ component (*** p<0.0001, ** p<0.001, * p<0.05).

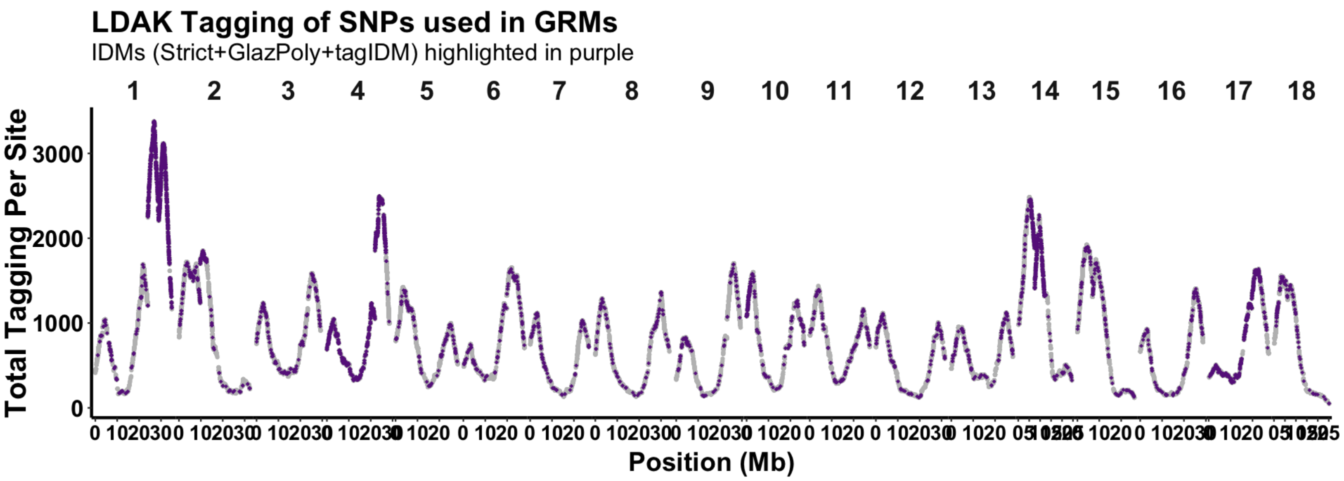


**Fig. S9. LDAK Tagging of SNPs used in GRMs.** Plot showing the LDAK score measuring the total amount of LD for each SNP (y-axis; column 4 from LDAK output) vs. its chromosomal coordinates (megabases, Mb). IDM SNPs (including tag-IDM) are highlighted in purple, non-IDM SNP are gray.

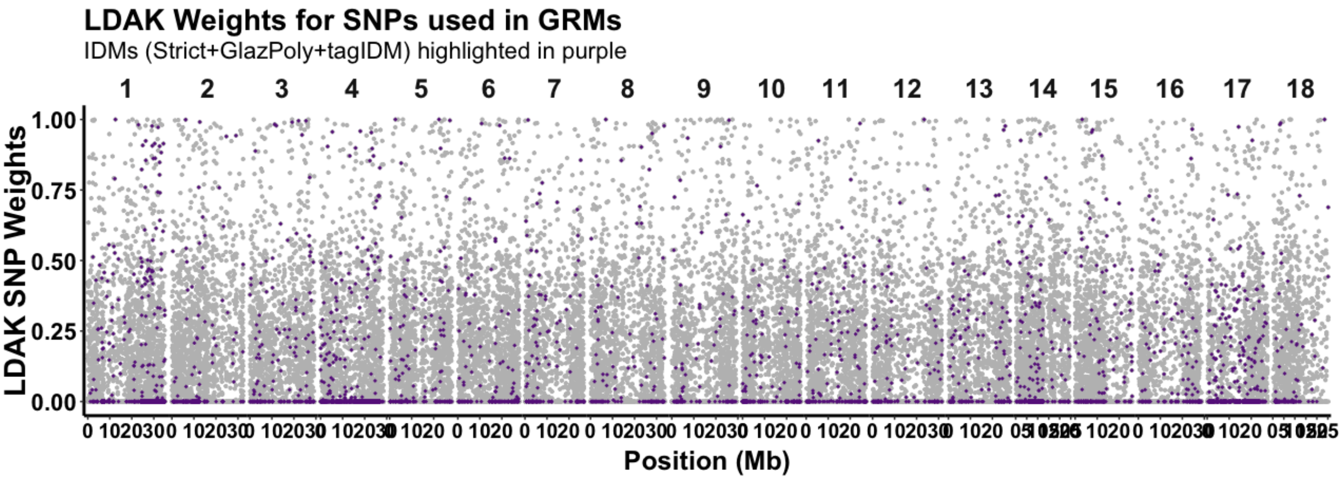


**Fig. S10. LDAK Weights used for weighting SNPs contribution to GRMs.** Plot showing the LDAK weight for each SNP (y-axis) vs. its chromosomal coordinates (megabases, Mb). The weight adjusts the contribution of each SNP to the kinships measured in an LD-adjusted genomic relationship matrix (GRM). IDM SNPs (including tag-IDM) are highlighted in purple, non-IDM SNP are gray.

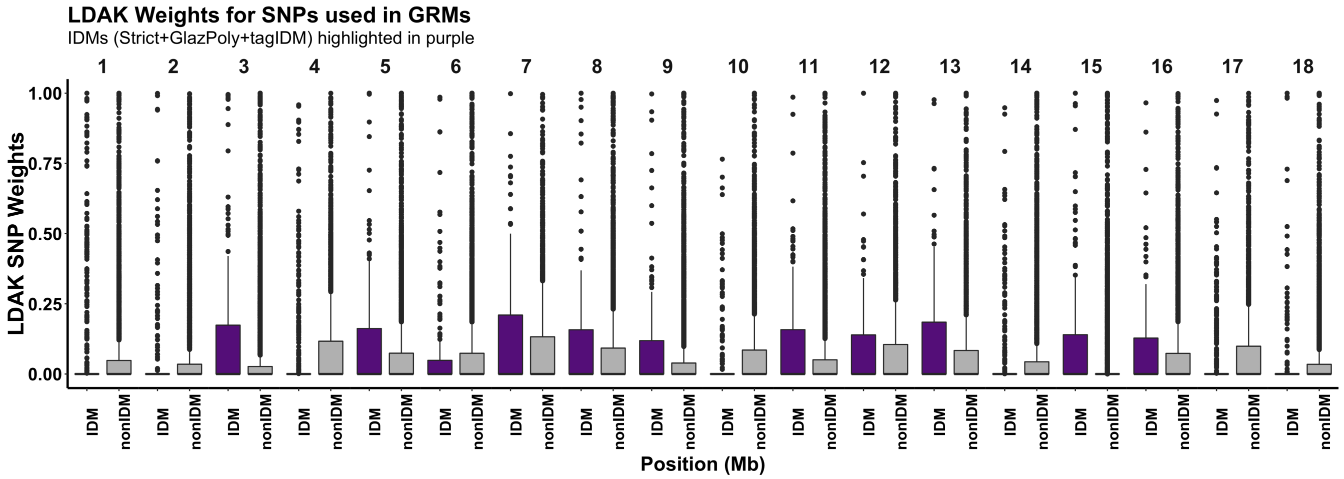


**Fig. S11. LDAK Weights for each chromosome, comparing IDM to non-IDM SNPs.** Boxplot shows the LDAK weight for each SNP (y-axis) vs. its status as an IDM SNPs (including tag-IDM) or not (x-axis) for each chromosome (horizontal panels). The LDAK weight adjusts the contribution of each SNP to the kinships measured in an LD-adjusted genomic relationship matrix (GRM).

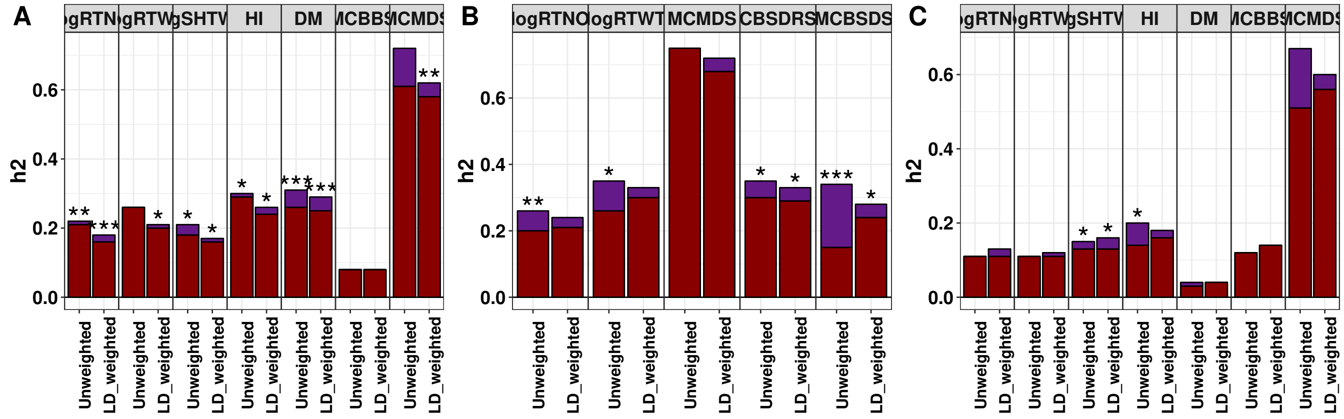


**Fig. S12. Heritability attributable to introgressions with vs. without LD-adjustment.** The heritability (y-axis) of introgression regions for each trait (horizontal panels) is shown for each breeding program (A = IITA. B = NaCRRI. C = NRCRI.). Results for two models are shown (x-axis): one where the genomic relationship matrix (GRM) was LD-adjusted using the LDAK method, and the other without LD-adjustment. In either case, heritability was estimated from partitioned genomic mixed-models and the portion of heritability attributable to introgression regions (purple) vs. the rest of the genome (dark red) is shown. Stars atop bars represent the level of significance in a likelihood ratio test for the significance of the introgression-component (*** p<0.0001, ** p<0.001, * p<0.05).

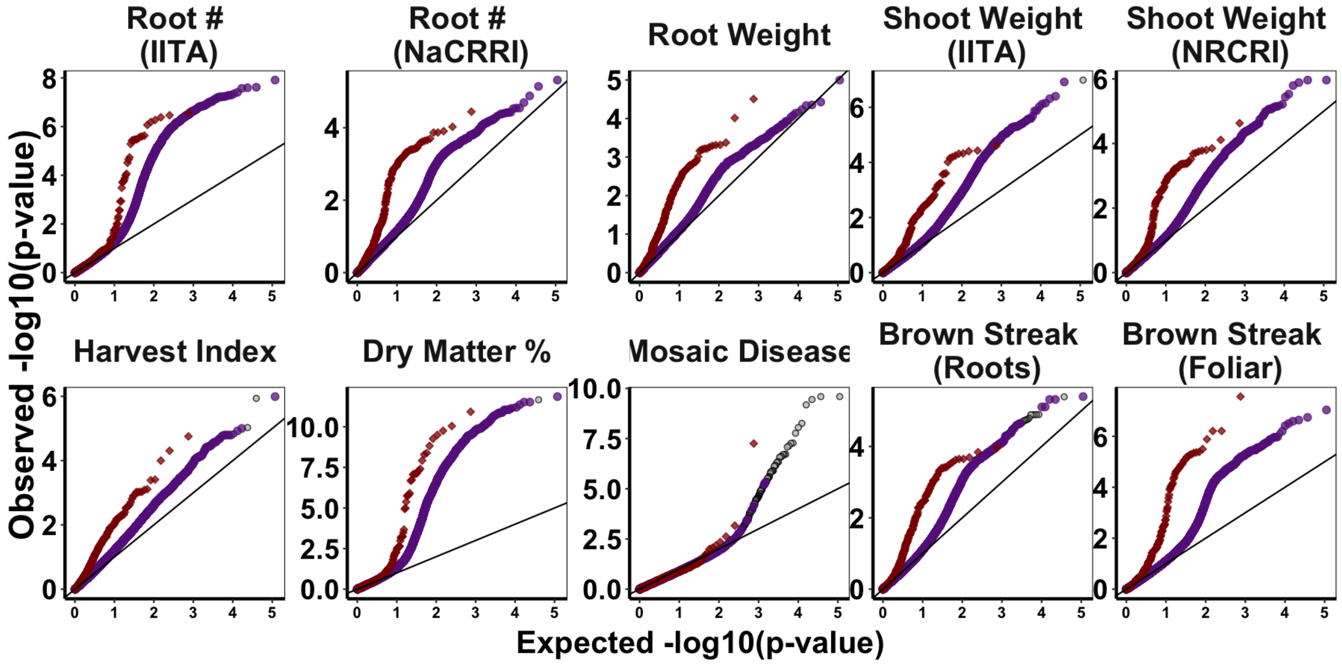


**Fig. S13. Quantile-quantile plots for each introgression-trait associations analysis.** Each panel shows QQ-plots for two-types of GWAS for one of 10 Trait-Institute analyses that had bonferroni-significant evidence of introgression-related associations. Two mixed-linear model association analyses are shown, overlayed. In the first, GWAS was conducted on IDM (purple) and non-IDM (gray) SNP markers. For the second, GWAS was done using using the mean *M. glaziovii* allele dosages in 250Kb windows (*DoseGlaz*; dark red diamonds).

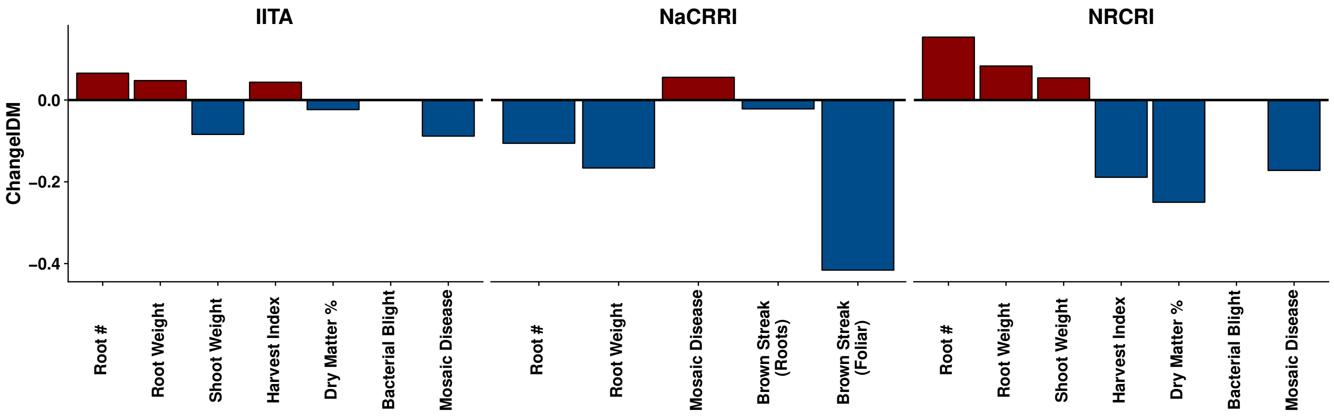


**Fig. S14. The change in the proportion of the heritability attributable to introgressions due to LD adjustment of the kinship matrix.** For each trait (x-axis) in each breeding program (horizontal panels), the proportion of the total heritability that was due to the introgression component (*Prop_h2IDM*) was computed for two models: one where the genomic relationship matrix (GRM) was LD adjusted using the LDAK method, and the other without LD adjustment. The difference between *Prop_h2IDM* between LD adjusted and un-adjusted estimates (*ChangeIDM*) is plotted on the y-axis. Positive values of *ChangeIDM* (dark red) indicate that *Prop_h2IDM* was larger for the LD adjusted model, and vice versa for negative values (blue).

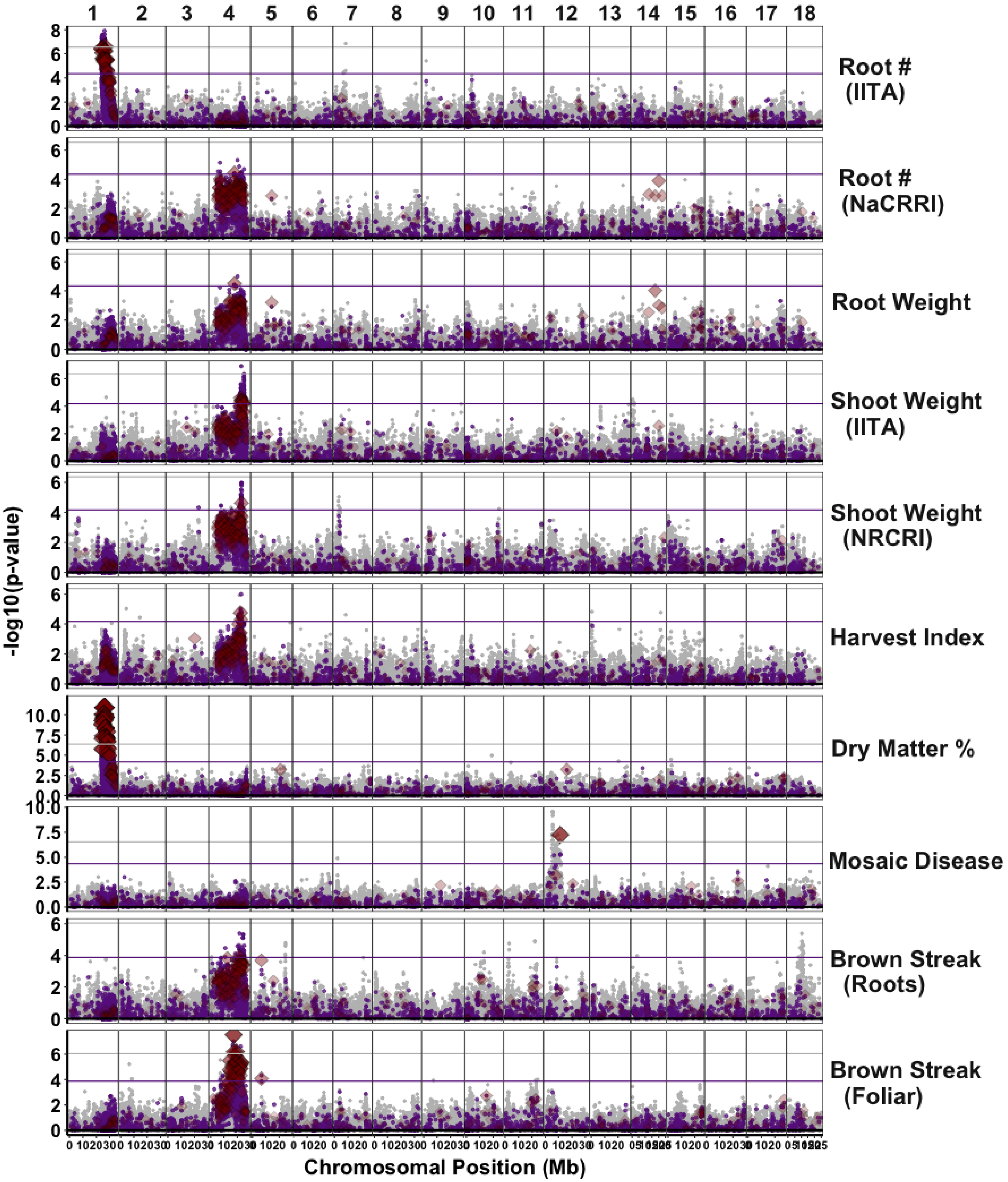


**Fig. S15. Genome-wide plot of significant introgression-trait associations.** Manhattan plots summarizing genome-wide associations is presented for 10 Trait-Institute analyses. Two mixed-linear model association analyses are shown, overlayed. In the first, GWAS was conducted on IDM (purple) and non-IDM (gray) SNP markers. For the second, GWAS was 250Kb-window mean *M. glaziovii* allele dosages (*DoseGlaz*; dark red diamonds). The size and alpha of *DoseGlaz* results are scaled with their $-log_{10}(p - value)$. The horizontal lines represent the bonferroni-significance threshold for the *DoseGlaz* (purple line) and SNP GWAS (gray line).

**Additional data table S1 (SI_Dataset.xlsx)**

**Introgression Diagnostic Markers.** For each marker in the dataset, indicates chromosome, position, genetic map coordinates (cM) and whether SNP is an IDM (Strict, GlazPoly or tagIDM) or not (nonIDM). Also records *M. glaziovii* allele (GlazAllele) at IDMs.

**Additional data table S2 (SI_Dataset.xlsx)**

**Raw per individual introgression summary.** Summary statistics for each cassava clone. Per individual metrics: M. glaziovii (PropMg) and deleterious (PropDeleterious) allele frequencies, M. glaziovii (PropHomMg) and deleterious (PropHomDeleterious) homozygote frequencies. Regions: per indidual summaries were calcluated either across all SNPs (GenomeWide) or only for SNPs in the introgression regions on chrs 1 ($> 25Mb$) and 4 (5-25Mb).

**Additional data table S3 (SI_Dataset.xlsx)**

**Population level summary.** Summary statistics for each cassava population. Per population statistics (min, mean, max) for each of the following per individual metrics: M. glaziovii (PropMg) and deleterious (PropDeleterious) allele frequencies, M. glaziovii (PropHomMg) and deleterious (PropHomDeleterious) homozygote frequencies. Regions: per population summary of per individual metrics calculated either across all SNPs (GenomeWide) or only for SNPs in the introgression regions on chrs 1 ($> 25Mb$) and 4 (5-25Mb).

**Additional data table S4 (SI_Dataset.xlsx)**

**Correlation of Kinships.** The correlation between the lower triangular off-diagonals of pairs of kinship matrices calculated with different combinations of markers according to the variables Matrix1 and Matrix2. Different criteria (LDscore, maxLD or None) with varying thresholds of the corresponding criterion for inclusion of SNPs in the kinship matrices.

**Additional data table S5 (SI_Dataset.xlsx)**

**Field Trial Summary.** Summary for each trait in each field trial that we analyzed. In addition to location, year, trial and trial type, meta information about the numbers of observations, clones and replications are provided.

**Additional data table S6 (SI_Dataset.xlsx)**

**Per-trial Model Summary.** For each trait-trial combination, we fit several models, as described in the text. Likelihood ratio tests and AIC-based model comparisons are given. Whether or not a trial passed three preliminary filters is also indicated.

**Additional data table S7 (SI_Dataset.xlsx)**

**Random partitions.** For each SNP, gives the status as either IDM or nonIDM in each of three random samples.

**Additional data table S8 (SI_Dataset.xlsx)**

**Cor. of randomly partitioned GRMs.** The correlation between lower triangular off-diagonals of pairs of the IDM-sized set of SNPs and the non-IDM sized set. SNP sets were partitioned either at random or based on true IDM-status.

**Additional data table S9 (SI_Dataset.xlsx)**

**LDAK Weights.** Weights and related raw output from LDAK.

**Additional data table S10 (SI_Dataset.xlsx)**

**Per-trial Per-model LogLik, AIC, VarComps, Etc.**

**Additional data table S11 (SI_Dataset.xlsx)**

**Summary of curated multi-trial data.** Indicates the sample size (Nobs) along with numbers of clones (Nclone), trials (Ntrial), replications (Ntrial_rep) and a summary of the distribution of observations per clone (nPerGeno suffix).

**Additional data table S12 (SI_Dataset.xlsx)**

**Summary of multi-trial analyses.** Gives the p-value for the likelihood ratio test for the significance of the IDM vs. non-IDM partition of genetic variance (LRT_partition). Whether the ALL or PARTITIONED model is better or the same is given based on which had a smaller AIC and whether there were more than 2 units AIC difference respectively (PARTvsALL). AIC comparison for the PARTITIONED vs. the IDMnull models are also given (PARTvsIDMnull). Heritability of the IDM and nonIDM parts (h2_IDM and h2_nonIDM) are also shown. Results are given for both IDM-based and random partitions, with and without LDAK LD weighting of the kinship matrices.

**Additional data table S13 (SI_Dataset.xlsx)**

**Multi-trial Per-model LogLik, AIC, VarComps, Etc.** Results are given for both IDM-based and random partitions, with and without LDAK LD weighting of the kinship matrices.

**Additional data table S14 (SI_Dataset.xlsx)**

**Compare to random.** For each random partition of the genome, compare it to the IDM-based partition. Compare AIC and the p-value from the likelihood ratio test on the significance of the variance component associated with the IDM-sized set of SNPs.

**Additional data table S15 (SI_Dataset.xlsx)**
**LDAK vs. A.mat GRMs.** Correlate the lower-triangular off-diagonals and the diagonals of the LDAK weighted and unweighted kinship matrices. Compare LDAK to A.mat (unweighted) kinship matrices calculated with either ALL, IDM, or nonIDM SNPs.

**Additional data table S16 (SI_Dataset.xlsx)**
**LDAK vs. A.mat cor(IDM,nonID).** Correlate the lower-triangular off-diagonals and the diagonals of the IDM vs. nonIDM (IDMvREST) kinship matrices. Compare A.mat (unweighted) to LDAK adjusted (weighted).

**Additional data table S17 (SI_Dataset.xlsx)**
**Compare to LD-adjusted.** For each Trait-Institute multi-trial dataset, compare the model results for the LDAK weighted and unweighted kinship matrices. Compare on the basis of AIC and change in the variance component associated with IDMs.

**Additional data table S18 (SI_Dataset.xlsx)**
**Summary of GWAS results.** For each Trait and type of GWAS summarize the per chromosome bonferroni-significant results. Indicates the number (Nsig), min and max position (minPos, maxPos) and the *M. glaziovii*-allele effect (Mean, Min, Max).

**Additional data table S19 (SI_Dataset.xlsx)**
**Cross-validation results.** Prediction accuracy from each rep of five-fold cross-validation conducted on each Trait-Institute-SampleMethod-Model and calculated for each variance component where relevant.

**Additional data table S20 (SI_Dataset.xlsx)**
**Summary of cross-val. Results.** Cross-validation accuracy results summarized for each Trait-Institute-SampleMethod. The mean (across reps) differences in accuracy between models is given.

**Additional data table S21 (SI_Dataset.xlsx)**
**Summary of genetic distance per megabase.** The cumulative genetic distance in centimorgans for each one megabase region along the genome, highling major introgression regions (distal 25Mb of chr. 1 and from 5-25Mb on chr. 4).
